# Different ecological processes drive the assembly of dominant and rare root-associated bacteria in a promiscuous legume

**DOI:** 10.1101/2020.01.13.900290

**Authors:** Josep Ramoneda, Jaco Le Roux, Emmanuel Frossard, Beat Frey, Hannes Andres Gamper

## Abstract

Understanding how plant-associated microbial communities assemble and the roles they play in plant performance are major goals in microbial ecology. For nitrogen-fixing rhizobia, assemblages are mostly determined by filtering by the host as well as abiotic soil conditions. However, for legumes adapted to highly variable environments and nutrient-poor soils, filtering out rhizobial partners may not be an effective strategy to ensure symbiotic benefits. As a consequence, this can lead to net increases in rhizobial diversity driven by stochastic (neutral) assembly processes. Here, we addressed whether symbiotic promiscuity of rooibos (*Aspalathus linearis* Burm. Dahlgren), reflects increases in rhizobial diversity that are independent of the environmental conditions, following a neutral assembly. We performed a common garden experiment to assess whether root system size and location- and habitat-specific rhizobial propagule pools of rooibos affected rhizobial community diversity and composition. We found a dominance of drift in driving taxonomic turnover in the root nodule communities, but operating at different scales in the dominant *Mesorhizobium* symbionts and the rest of bacterial taxa. Specifically, drift triggered differentiation between the core rhizobial symbionts of rooibos due to dispersal limitation on a regional scale, whereas it dominated the assembly of non-dominant rhizobial taxa at the root level. This suggests the existence of a significant neutral component in rhizobial community assembly when selectivity by the host plant is low. We conclude that in this promiscuous legume neutral processes govern bacterial community root nodule community assembly, but that these operate at different scales in dominant and rare rhizobial symbionts of the plant.

## Introduction

Exploring ways to manage entire microbial communities with beneficial functions on plant nutrition and health (i.e. functional microbiomes) is a recognized research priority (Lemanceau *et al.*, 2017; Toju *et al.*, 2018). Under field conditions, microbial community assembly from the plant’s surrounding environment defines which taxa and potential functions will assist plant health and growth. Therefore, we need a better understanding of how microbial communities assemble in plants, and the roles microbial diversity can play in plant performance, both from ecological and evolutionary standpoints.

Legumes are highly reliant on microbial services and key to ecosystem productivity and agricultural soil fertility (Graham and Vance, 2003). This is because most of them associate with rhizobia from the alpha- and β-Proteobacteria, as well as other microbes that improve growth and alleviate stress impact. Rhizobia fix atmospheric nitrogen (N) in structures called root nodules. Rhizobia thrive as saprobes in soil and the rhizosphere and vastly multiply when in association with their legume hosts, within root structures called root nodules. The plant controls the symbiosis functioning by selectively allocating carbohydrates and other key resources for the endosymbiotic and functionally differentiated rhizobia (Denison and Kiers, 2011).

Surprisingly little is known about rhizobial community assembly in legume roots (Barrett *et al.*, 2015; Sprent *et al.*, 2017; Burghardt *et al.*, 2019). Until most recently, community assembly in rhizobia has received little attention because of presumed low functional diversity, genetically homogeneous rhizobial populations in root nodules, and lack of analytical technology (Burghardt *et al.*, 2019). At the local scale, symbiotic rhizobial community assembly is driven by soil abiotic factors (e.g. pH, humidity, soil texture), the nutritional status of their host plant (Denison and Kiers, 2011; Bever 2015), host plant occurrence and abundance (Dension and Kiers 2011), predators and parasites (Burghardt 2019). Additionally, at the landscape scale differences in population sizes of rhizobia can lead to evolution of geographical variants and ultimately speciation (Bissett *et al.*, 2010; Sprent *et al.*, 2017). Thus, site and host plant adaptation, along with differential propagation of the symbionts, determine the composition of available rhizobial propagules.

As for most symbiotic interactions, the assembly of rhizobial communities in legume roots has both deterministic and stochastic components (Nemergut *et al.*, 2013). These components dominate in so-called trait-based and neutral assembly mechanisms, respectively.

Deterministic assembly is driven by plant-microbe compatibility and adaptation, in which trait differences are decisive. On the other hand, stochastic assembly is reliant on the chance of partner encounters, infection pressure, infection priority or propagule dispersal (Nemergut *et al.*, 2013; Le Roux *et al.*, 2017). Legume-rhizobium partners associate *de novo* at the onset of each new plant generation, but also every time when new root nodules and new roots are formed, and during root growth. These events provide chances for both trait-based and neutral assembly processes to operate. However, their relative importance in legume-rhizobium associations are far from being understood, particularly under agricultural settings.

For legumes thriving in nutrient-poor environments, where extreme N limitation is the norm, it can be advantageous to associate with any, even functionally inferior rhizobia (Kiers *et al.*, 2003). Indeed, legumes in these environments seem to lack several of the mechanisms for symbiont sanctioning (cf. Australian acacias or legumes in the Cape Floristic Region of South Africa: Lemaire *et al.*, 2015; Brink *et al.*, 2017; Vuong *et al.*, 2017; Le Roux *et al.*, 2018). Such legumes associate with higher rhizobial diversity, depending on factors such as drought, plant health, root system size, or plant age (Jiao *et al.*, 2015; Wagner *et al.*, 2016; Dinnage *et al.*, 2019). In the absence of legume control mechanisms, propagule pressure, together with infection priority, may render abundant rhizobium taxa monodominant in root nodules, especially during seedling establishment. Over time, the probability of root-rhizobium encounters may rise as older legume plants associate with increasing numbers of different rhizobia (Dinnage *et al.*, 2019). This may lead to root size-dependent assembly of rhizobial root symbionts, which is apparently neutral, even though infection and maintenance of symbioses at fine (i.e. root) scales may still be trait based (Yu *et al.*, 2018).

To better understand the links between root nodule symbiont diversity and neutral vs non-neutral community assembly processes, we studied rhizobial communities associated with rooibos (*Aspalathus linearis* Burm. Dahlgren), an economically important crop. The species is endemic to the acidic and extremely nutrient-poor soils of the semi-arid South African Fynbos vegetation, and its agricultural production is under threat by climate change and desertification (Malgas *et al.*, 2010; Lötter *et al.*, 2014). Rooibos belongs to the diverse genus *Aspalathus*, which has radiated exclusively in the hyperdiverse Core Cape Subregion (CCR, previously Cape Floristic Region, and now forming part of the Greater Cape Floristic Region) in close association with a wide diversity of rhizobial symbionts (Hassen *et al.*, 2012).

Given the extreme nutrient-poor conditions under which rooibos evolved, we hypothesized that the assembly of the rhizobial root nodule communities of the species would follow predominantly stochastic rather than deterministic trajectories. Neutral assembly was expected to lead to increased rhizobial diversity when exposed to more diverse inocula and when having larger root systems (i.e. under conditions of increased likelihood of encounter and infection). Using a dual marker approach, we described both symbiotic and non-symbiotic rhizobial communities, and manipulated soil nutrient content and soil inoculum diversity. By using soils from a range of geographical locations, and covering a range of plants with different root sizes, we were able to link assembly processes to regional and root spatial scales.

## Materials and Methods

### Experimental design and setup

To examine the effects of fertilization and soil mixing on the assembly of root-nodulating rhizobia in rooibos, a cross-factorial pot experiment was setup. It was designed with the factors i) ‘location’ with five levels, represented by five farm origins of the employed soil, ii) ‘soil origin’ with three levels, made up of soil from adjacent cultivated and uncultivated land and a 1:1 (v:v) volumetric mixture of those two, and iii) ‘fertilization’ with two levels, which were either addition of sheep dung or not. The experimental units were pots with one plant individual that were randomly arranged with respect to all 30 resulting experimental treatments (5 locations × 3 soil origins × 2 fertilization treatments) in ten spatial blocks, representing ten replicates, totalling in 300 experimental units represented by individual pots with one single rooibos seedling.

The soils used for the experiment were collected down to a depth of 30 cm from beneath cultivated and wild rooibos plants not more than 100 m apart from each other, and locations that corresponded with different farms in the Suid Bokkeveld (Western Cape, South Africa). The locations were located 5.5-33 km apart from each other. The experiment was initially setup in a polytunnel and after five months transferred to an outdoor shade house at the Agricultural Research Council Infruitec’s research station in Stellenbosch, South Africa.

Two-liter pots were filled with soil from either cultivated or uncultivated land or a 1:1 volumetric mixture of those. The fertilization treatment consisted in the addition of 7.5 g air-dry dung from a nighttime enclosure (Afrikaans ‘kraal’) of sheep on one of the farms where soils were collected. The sheep dung increased the total N content in the soil of the pots by 30% and the total P content by 50% (see Supplementary Table 1 for the detailed mineral nutrient concentrations of the sheep dung).

Three sulfuric acid-scarified seeds of the sole cultivated rooibos variety *Nortier* were buried about 1 cm below the soil surface in a line perpendicular to that defined by the two patches of sheep dung to avoid direct root-dung contact initially and hence prevent any possible nutrient toxicity. In the last two months post germination, the strongest seedling was kept and any remaining seedling removed to finally have one plant per pot. Other plants growing from the seedbank or from root fragments were periodically removed throughout the duration of the experiment. The experiment ran from the beginning of August 2016 to the middle of April 2017, corresponding to the period between early spring and late autumn under the South African Mediterranean climate.

### Harvest and sample preservation for the various analyses

At harvest, individual root systems were well rinsed with pressurized water. Stems, leaves, and roots were separated and the number of root nodules and cluster roots counted. The dry weight of these separate plant fractions was determined after air-drying at 50°C for two days, except for the root nodules which were removed prior to drying. Root nodules were initially air-dried for 1–3 hours before being stored in silica gel. After drying, leaves were removed from stems and branches and weighed separately. The maximum diameter of the taproot was recorded with a caliper. Dried leaves were finely ground for nutrient analyses in wolfram carbide cups on a TissueLyser II swing mill (Qiagen, Hombrechtikon, Switzerland) for 60 s at 28 Hz.

### Mineral nutrient and δ^15^N signature analysis

The foliar carbon (C) and nitrogen (N) concentrations and δ^15^N isotopic signature were determined from 4 mg subsamples of milled leaf material on an ion ratio mass spectrometer (Delta V Advantage, ThermoFisher Scientific, Waltham, MA, USA). The other macro- and micronutrients (P, K, Ca, Mg, Mn, Fe) were quantified in 200 mg of ground leaf material, which was digested in 2 ml HNO_3_(69%) and 2 ml deionized H_2_O in a microwave liquid digestion system MLS Turbowave (MWS GmbH, Heerbrugg, Switzerland) and analyzed by Inductively Coupled Plasma – Optical Emission Spectrometry (ICP-OES) on a ICPE-9800 apparatus (Shimadzu, Kyoto, Japan).

The soil pH in 0.01M CaCl_2_ was measured in triplicate technical replicates. The ^15^N isotopic signature of triplicate 3 mg subsamples of finely ground sheep dung and macronutrients of finely ground 200 mg subsamples of the soils and sheep dung were measured after liquid extraction with 4 ml of HNO_3_(69%) and 10 ml deionized H_2_O as described above for the plant tissue. All plant nutrient analyses were carried out in the laboratory facilities of the Group of Plant Nutrition at ETH Zurich (Eschikon, Switzerland). The physico-chemical characteristics of the soils were determined by the ISO 17025:2005-certified Bemlab laboratories (Somerset West, South Africa).

### DNA extraction, partial gene amplification and sequencing

The root nodules of each individual plant were pooled and milled using a 7 mm diameter glass bead in a 2 ml Eppendorf tube in a TissueLyser II (Qiagen, Hilden Germany). The total DNA was then extracted from 25 mg of nodule powder using the NucleoSpin Plant II Kit (Macherey-Nagel, Düren, Germany) according to the manufacturer’s protocol with the sodium dodecyl sulfate-containing lysis buffer PL2 and elution in 100 μl 5mM Tris-EDTA (TE, pH 8.5) elution buffer (PE).

A 817 bp-long fragment of the DNA gyrase B (*gyrB*) gene was amplified using fusion primers with the *gyrB*343F gene-specific primer sequence (Martens *et al.*, 2008) and PacBio’s 30mer *universal sequence* as a tag at the 5’-end (5’-*GCAGTCGAACATGTAGCTGACTCAGGTCAC*TTCGACCAGAAYTCCTAYAAGG-3’) and the gene-specific primer *gyrB*1043 (Martens *et al.*, 2008) and PacBio’s 30mer *universal sequence* as a tag at the 5’-end (5’-*TGGATCACTTGTGCAAGCATCACATCGTAG*AGCTTGTCCTTSGTCTGCG-3’). Both primers were amino-blocked at their 5’-end (5’ NH4-C6) to prevent their ligation during library preparation and hence sequencing without a sample-specific barcode. As an easy to align and variable bacterial housekeeping gene, *gyrB* has proven useful as a phylotaxonomic marker to analyze legume root nodule inhabiting bacteria (e.g. Beukes *et al.*, 2016). PCR reactions were performed in triplicates, using a final reaction volume of 12.5 μl. Each reaction contained 4.25 μl PCR-grade water, 6.25 μl Multiplex PCR Master Mix (QIAGEN Multiplex PCR Plus Kit), 0.5 μl of each of the 10 μM primers and 1 μl of the template DNA. The amplification reactions were run in a DNA Engine Peltier Thermal Cyclers (Bio-Rad, Hercules, CA, USA), using the following conditions: initial denaturation at 95°C for 5 min followed by 30 cycles of 95°C for 30 s, primer annealing at 58°C for 2 min and primer extension at 72°C for 45 s, followed by a final extension phase at 68°C for 10 min. Reactions were kept at 10°C until storage at −20°C. The three replicated PCR reactions of each sample were pooled, and the amplicons purified with home-made Solid Phase Reversible Immobilization (SPRI) magnetic beads at a ratio of 0.7 times the reaction volume (Genetic Diversity Center, ETH Zürich, Switzerland).

The amplicons of the different samples were uniquely barcoded with PacBio’s 96 Barcoded Universal F/R Primers (Pacific Biosciences, Menlo Park, CA, USA) in a second PCR step for later sample pooling and library construction for Single Molecule Real-Time (SMRT) circular consensus sequencing (ccs). These barcoding primers consist of the complementary 30-mer primer sequence to the universal sequences and contain at their 5’-ends 16-mer unique barcodes. The PCR reaction mix of the barcoding PCR contained 12.25 μl PCR-grade water, 5 μl 5x Q5 reaction buffer (New England Biolabs), 2.5 μl dNTPs (2mM), 2.5 μl of each of the index primers (10 μM), 0.25 μl of Q5 polymerase (New England Biolabs, Ipswich, MA, USA), and 2.5 μl of the bead-purified amplicons diluted to 1 ng μl^−1^ of the first PCR as template. The following thermal cycling program was used: initial denaturation at 98°C for 30 s, followed by 10 cycles of denaturation at 98°C for 10 s, primer annealing at 71°C for 20 s and primer extension at 72°C for 60 s, with a final further extension at 72°C for 2 min. The barcoded amplicons were again purified using the same SPRI magnetic bead-based protocol. The concentrations after purification were measured with the Qubit assay (Thermo Fisher Scientific Inc., Waltham, MA, USA) on a Spark 10M Multimode Plate Reader (Tecan, Männedorf, Switzerland).

For construction of sequencing libraries, all bead-purified barcoded amplicons were diluted to a concentration of 10 ng μl^−1^ and multiplexed in pools of 96 samples, whose integrity was checked with an Agilent 2200 Tape Station (Agilent, Santa Clara, CA, USA). Three sequencing libraries were prepared with the SMRTbell™ Template Prep Kit 1.0‐SPv3 (Pacific Biosciences) chemistry by the Functional Genomics Center Zurich (University of Zurich, Zurich, Switzerland) and SMRT circular consensus sequenced on individual SMRT cells on a Sequel PacBio sequencer. The average output was 3726 sequence reads per sample, covered by on average 55 and a minimum of 5 polymerase passes, which guarantees a minimum predicted sequencing accuracy of 0.999.

A 455 bp-long fragment of the nodulation gene *nodA*, encoding an N-acyltransferase of the rhizobial *nodA*BC operon, was amplified with the gene-specific primers *NodA*univF145u (5’-TGGGCSGGNGCNAGRCCBGA-3’) and *NodA*Rbrad (5’-TCACARCTCKGGCCCGTTCCG-3’) (Moulin *et al.*, 2001). The PCR amplifications were performed in triplicates in final reaction volumes of 12.5 μl. Each reaction contained 3.75 μl PCR-grade water, 6.25 μl Multiplex PCR Master Mix (QIAGEN), 0.75 μl of each of the primers at a concentration of 10 μM and 1 μl of the template DNA. The amplification reactions were run in DNA Engine Peltier Thermal Cyclers (Bio-Rad), using the following conditions: initial denaturation at 95°C for 5 min followed by 27 cycles of denaturation at 95°C for 30 s, primer annealing at 69°C for 90 s and primer extension at 72°C for 35 seconds, with a final further extension at 68°C for 10 min. The reaction products were kept at 10°C until storage at −20°C. The three amplification product replicates of each root nodule extract were pooled and purified with home-made SPRI beads at a ratio of 0.8 times the reaction volume. After dilution to 1 ng μl^−1^ unique Nextera XT sample indices (Illumina, San Diego, CA, USA) were added by a second PCR in a reaction volume of 25 μl with 12.5 μl KAPA HiFi Hot Star Ready Mix (KAPA Biosystems Inc., Wilmington, MA, USA), 5 μl PCR-grade water, 2.5 μl of each of the index primers at a concentration of 10◻M and 2.5 μl of the purified amplicons as template. The thermal cycling program was the following: initial denaturation at 95°C for 3 min, 8 cycles of denaturation at 95°C, primer annealing at 55°C and primer extension at 72°C for 30 s and a final further extension at 72°C for 5 min. The indexed PCR products were bead-purified again, using the same protocol as for the initial amplicons and quantified, using the Qubit assay (Thermo Fisher Scientific Inc.) on the Spark 10M Multimode Plate Reader (Tecan). The amplicon size and integrity was verified on the Agilent 2200 Tape Station (Agilent, USA) before dilution to 4 ng ◻l^−1^ for equimolar pooling of all 266 samples for library preparation. The amplicons were 2×300 bp paired-end sequenced, using the Illumina MiSeq v2 chemistry and PhiX as internal standard at a concentration of 48.99% on an Illumina MiSeq sequencer at the Genetic Diversity Center of ETH Zürich, Switzerland.

### Bioinformatics of sequencing data

For the *gyrB* ccs reads, the SMRT link software (Pacific Biosciences) was used to remove the bell adaptor, barcode and primer sequences, filter the reads for a size range of 700-1000 bp (expected size 817 bp) and demultiplex the sequences of different samples. Read quality assessment and filtering was done in PRINSEQ-lite (Schmieder and Edwards, 2011). All sequences across the entire dataset were demultiplexed in USEARCH, using the function *fastx_uniques* to obtain only the unique sequences for subsequent further quality measures, similarity clustering and phylotaxonomic assignment. The unique sequences were error-corrected and chimera-filtered in UNOISE3, which generated zero-radius operational taxonomic units (ZOTUs). These are expected to represent the true biological sequences and hence possibly different bacterial strains. To further exclude possible variants due to PCR and sequencing errors, we clustered the ZOTUs at 99% nucleotide sequence identity, using the USEARCH function *cluster_smallmem*. The count table containing the information about the read numbers and thus ZOTU abundance was extracted, using the USEARCH function *otutab*. Taxonomic assignment and a chimera check were done in SINTAX (Edgar, 2016), using all available *gyrB* sequences from NCBI GenBank as the reference database (https://blast.ncbi.nlm.nih.gov/Blast.cgi).

For the *nodA* paired-end reads, seqtk (https://github.com/lh3/seqtk) was used for 3’-end trimming and FLASH (Magoč and Salzberg, 2011) to join the two ends. The priming sites were removed with USEARCH (v10.0.240) as described in Edgar (2011). PRINSEQ-lite (Schmieder and Edwards, 2011) was used to check the quality of the sequence dataset. The unique sequence reads were obtained with the dereplication command *fastx_uniques* in USEARCH and Illumina sequencing errors were corrected and chimeras removed in UNOISE3. ZOTUs were determined using the same approach as for the *gyrB*. The count table was generated, using the *otutab* command in USEARCH. The phylotaxonomic affiliations of the ZOTUs and a further chimera check were done in SINTAX (Edgar, 2016) in comparison to all *nodA* sequences on NCBI GenBank.

The relative abundances of the ZOTUs were intra- and extrapolated to the median number of reads per sample (1410 reads for *gyrB*, 8720 reads for *nodA*) after excluding four samples with less than 200 reads from the *nodA* dataset and twelve samples with less than 300 reads from the *gyrB* dataset, using the *iNEXT* package (v.2.0.19, Hsieh *et al.*, 2016) in R. *iNEXT* was also used to calculate the α-diversity (richness and Simpson’s diversity index). All further raw sequence data processing was done in the *phyloseq* package (v1.16.2, McMurdie *et al.*, 2013) of R. A total of 931,254 *nodA* and 161,155 *gyrB* error-filtered sequence reads were analysed. An indicator species analysis, using the function *multipatt* in the R package *indicspecies* (v1.7.6) was used to infer those ZOTUs whose occurrence and relative abundance was indicative for a certain experimental treatment or plant size class.

### Statistical analysis

We ran generalized linear mixed model analyses with location and block as random factors in the R package *lme4* (v1.1-21) to test the effects of Fertilization and Soil origin on the plant growth physiological, symbiotic and alpha diversity parameters. The exact statistical model used was: *y* ~ *Fertilization*Soil origin*, *random= ~1|Block/Location.* P-values for the results of the mixed effect models were obtained by running ANOVA on the model fits. Means were compared by post-hoc Tukey’s HSD tests at the confidence level of 95%.

β-diversity was analyzed using the relative ZOTU abundances. Analyses on the Hellinger (square-root) transformed abundances did not change the results qualitatively. The effects of the experimental treatments on the composition and structure of the rhizobial communities in the root nodules were visualized by non-metric multidimensional scaling (NMDS) of the Bray-Curtis dissimilarities in the package *vegan* (v2.5-5, Oksanen *et al.*, 2007) of R. The treatment effects were statistically tested by a permutational analysis of variance (PERMANOVA) with the *adonis* function in *vegan*.

Possible causative relationships between soil abiotic parameters and the bacterial communities associated with the root nodules were assessed using correlation analyses. This is because we lacked paired data between soil characteristics and microbial communities and plants (Supplementary Table 1), making statistical testing impossible. Since we had only soil data per Location and Soil origin, the β diversity of the two fertilization treatments and ten experimental replicates was summarized as the distances between the centroids of the Location (5) × Soil origin (3) factor level combinations (15) of a PCoA ordination. These distance values were then correlated to the Euclidean distances between soil parameter values using the *dist_between_centroids* function of the *usedist* package (v0.1.0) of R. Possible effects of differences in the rhizobial community composition and structure (Bray-Curtis dissimilarities) on plant performance (Euclidean distances) were assessed by means of Mantel tests, using the *vegan* function *mantel.test*.

As a way to measure the relative importance of neutral and trait-based assembly processes, we followed the analytical framework described by Stegen *et al.*, (2012, 2013). The approach is based on the comparison of the phylogenetic distance between closely related taxa across samples to a null distribution. In a first step, we assessed the strength of species turnover using the β-nearest taxon index (β-NTI). This index quantifies how much taxonomic turnover there is between samples, by comparison to a null distribution. This null distribution, which represents the absence of filters in defining taxon presence or absence, is calculated on randomized phylogenetic distances between samples by randomly shuffling taxon identities and abundances across the tips of a phylogeny. β-NTI values below −2 and above +2 indicate that taxa between two communities are more or less phylogenetically related than expected by chance, respectively. These underlie a role for homogenizing and variable selection respectively (i.e. either the same or different taxa are selected in different communities; Stegen *et al.*, 2015). β-NTI values between −2 and +2 indicate no effects of selection in driving differences between two given communities, leaving only dispersal, dispersal limitation, and drift as possible drivers. To identify these alternative processes, in a second step we calculated the Raup-Crick index, which is the departure from a null distribution of Bray-Curtis dissimilarities between probabilistically generated communities (RC_Bray_). This distribution is created by generating random communities drawn from a species pool composed by the same species composition, richness and absolute abundance as in the original dataset. The probability of these communities of containing particular taxa is proportional to the abundance and frequency of occurrence of those taxa both in the particular samples being compared and in the entire dataset. This index does not rely on phylogenetic distances because phylogenetic relatedness is considered irrelevant for the probability a species has of being dispersed or subject to ecological drift. RC_Bray_ values below −0.95 or above +0.95 indicate two communities have more or less taxa in common than expected by chance, respectively. These are indicative of homogenizing dispersal and dispersal limitation plus drift, respectively. A RC_Bray_ value between −0.95 and +0.95 indicates two communities have as many taxa in common as expected by chance, and is therefore indicative of ecological drift. The β-NTI and RC_Bray_ were calculated with 999 randomizations (Stegen *et al.*, 2012, 2013). We used the R package *picante* (v1.7) to calculate the Faith’s phylogenetic distance and its function *comdistnt* for calculation of the mean nearest taxon index (β-MNTD) needed for calculation of β-NTI.

## Results

### Rooibos biomass and nutritional response to fertilization and soil origin

The only effect on rooibos growth and nutrition came from fertilization, which doubled rooibos biomass (Figure 1; Table 1). Although plants grown in mixed soils had on average higher biomass, this effect was not significant (Table 1). Moreover, there was no interaction between the fertilization and soil origin treatments. The absence of fertilizer triggered a physiological response in the form of increased cluster root production (Figure 1). In line with this observation, foliar N:P ratios shifted from 26.22 ± 7.15 to 18.57 ± 5.32 when fertilizer was added (P<0.001). Fertilization stimulated root nodule formation by increasing total plant dry matter, including that of the root system. Importantly, the number of root nodules correlated positively with root system dry matter, regardless of the fertilization treatment, while the intensity of nodulation per root dry matter remained the same (Table 1; Supplementary Figure 1). All macro- and micronutrient concentrations in the leaves and roots were higher in the fertilized plants, except the Ca in the shoots and Fe in the roots (Supplementary Table 2

**Table 1.** Results of mixed model analyses on the effects of fertilization, soil origin and their interaction on plant biomass, nitrogen (N) and phosphorus (P) nutritional parameters as well as cluster root and root nodule formation of rooibos. The experimental factor ‘fertilization’ involved sheep dung addition to soil, and ‘soil origin’ comprises use of soils from five pairs of plantations and adjacent uncultivated populations of rooibos and a 1:1 (v:v) mixture of those. The five locations (farms) were treated as a random and nested factor within fertilization and soil origin. There were 10 pot replicates per experimental treatment.

| Response variable | Experimental factors |  |  |  |  |  |
| --- | --- | --- | --- | --- | --- | --- |
|  | Fertilization (df = 1) |  | Soil origin (df = 2) |  | Fertilization:Soil origin (df = 2) |  |
|  | F-value | P-value | F-value | P-value | F-value | P-value |
| Total plant dry biomass (g) | 72.87 | **<0.001** | 2.31 | 0.102 | 0.82 | 0.440 |
| Root dry biomass (g) | 60.43 | **<0.001** | 2.01 | 0.137 | 0.76 | 0.468 |
| Leaf N content (mg) | 109.24 | **<0.001** | 1.71 | 0.183 | 0.51 | 0.600 |
| Leaf P content (mg) | 181.82 | **<0.001** | 3.26 | **0.040** | 0.94 | 0.392 |
| Leaf N:P ratio | 105.88 | **<0.001** | 11.24 | **<0.001** | 0.22 | 0.300 |
| □ <sup>15</sup>N (‰) | 211.76 | **<0.001** | 0.34 | 0.714 | 1.18 | 0.308 |
| # Cluster roots | 31.72 | **<0.001** | 0.30 | 0.740 | 0.57 | 0.566 |
| # Nodules | 59.98 | **<0.001** | 3.46 | **0.033** | 2.09 | 0.126 |
| Nodule density (# nodules root dry biomass<sup>-1</sup>) | 0.20 | 0.653 | 13.14 | **<0.001** | 2.64 | 0.073 |

**Figure 1.**
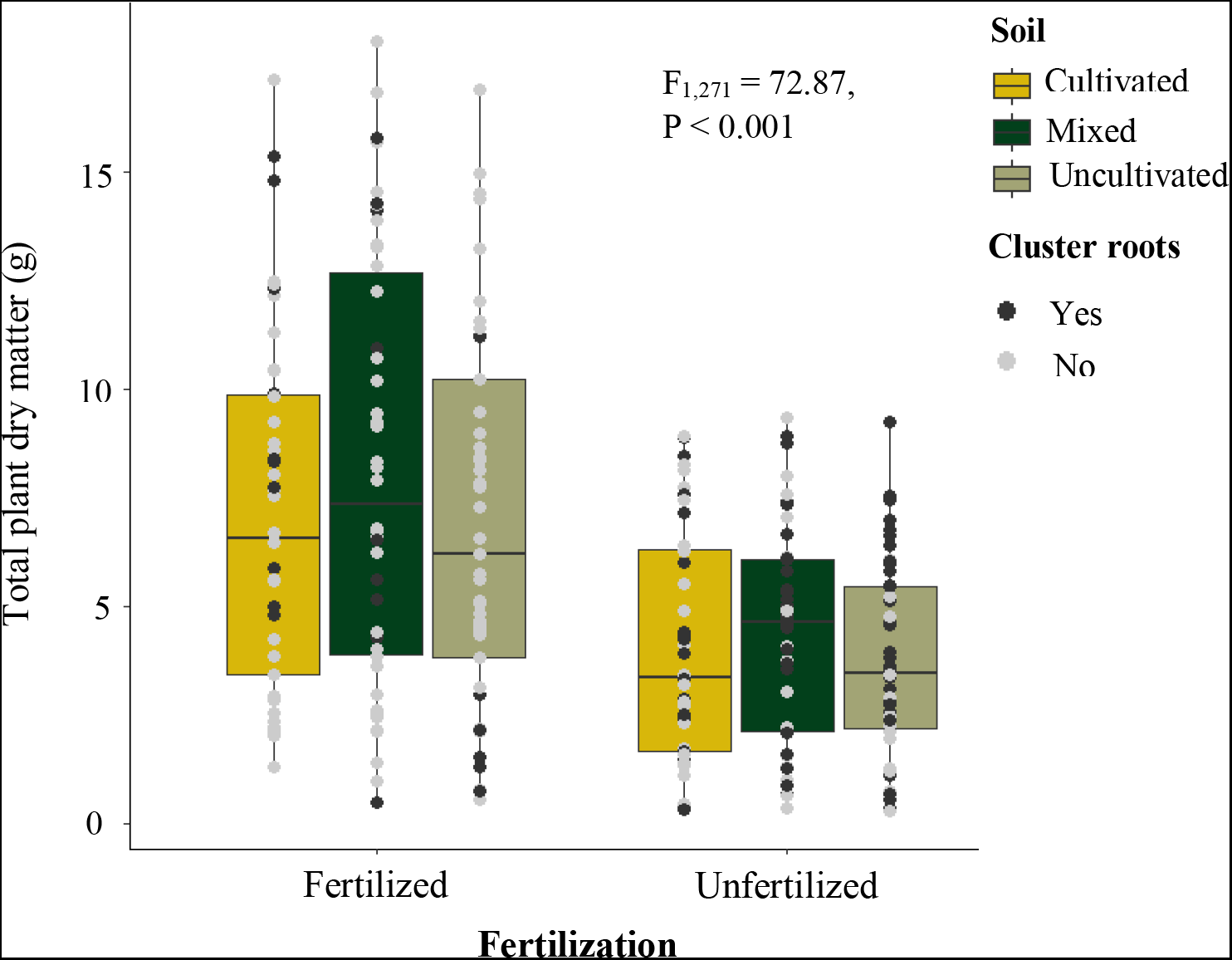
Total dry plant biomass of rooibos grown in pots for eight months as affected by fertilization with sheep dung and by soil from either plantations, wild populations, or a mixture of these. The soils were collected from five locations/farms with adjacent cultivated and uncultivated rooibos populations. Box plots show the median and first and third quartile and the whiskers 1.5 standard deviations. Black dots indicate data from plants with cluster roots and grey dots data from plants without cluster roots. The statistical summary reports on differences in plant biomass between the fertilization treatments (n=271).

### Alpha diversity of root nodule communities

A total of 93 different *nodA* and 355 *gyrB* ZOTUs were identified in the eight-month old rooibos seedling root nodules. These belonged exclusively to the genus *Mesorhizobium* based on *nodA*, while *gyrB* sequences also corresponded the genera *Rhizobium*, *Bradyrhizobium*, *Agrobacterium* and *Paraburkholderia*, the latter at extremely low abundances, apart from a few non-rhizobial taxa (Figure 2). All genera were present in all treatments, with no evident differences in their distribution with fertilization or soil origin. *Mesorhizobium* were by far the most dominant symbionts in rooibos root nodules, making up the most dominant ZOTU in all but one *gyrB* community. The most dominant ZOTUs associated with individual plants represented 93% and 86% of the average symbiont abundance in *nodA* and *gyrB* respectively. Plants grown in fertilized soils hosted a larger proportion of non-*Mesorhizobium* ZOTUs compared plants grown in non-fertilized soils (Figure 2), while soil origin also influenced their relative abundance. The effect of soil origin on relative abundances was especially strong in non-fertilized soils, with cultivated soils having a higher proportion of non-*Mesorhizobium* ZOTUs than the other soil treatments (Figure 2B).

**Figure 2.**
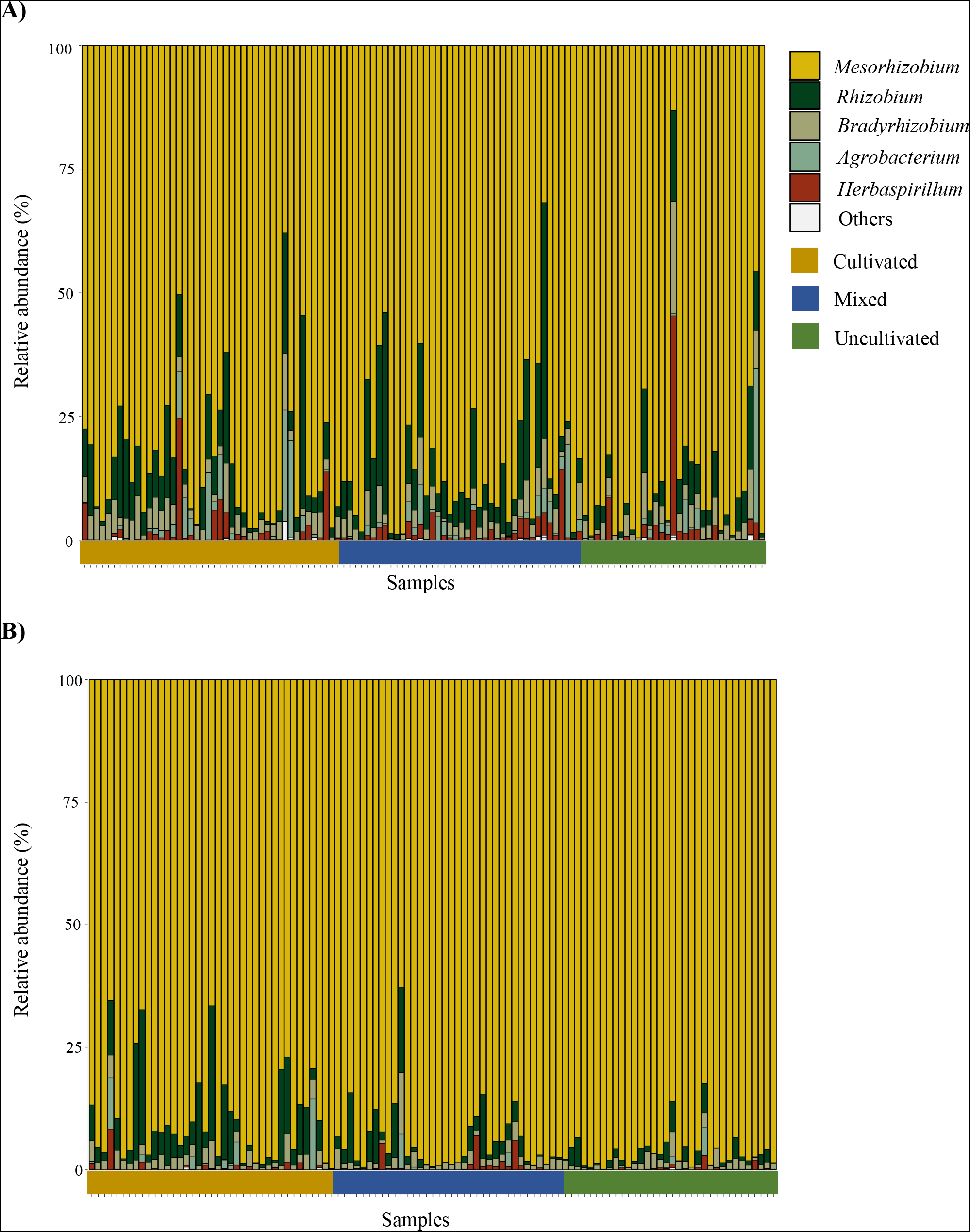
Relative abundances of different rhizobial genera from pools of root nodules of eight-month-old rooibos grown in soils from five plantations (cultivated) and adjacent wild (uncultivated) populations and a 1:1 (v:v) mixture of soil pairs (mixed). Soils were either fertilized with sheep dung (A) or not (B). The analysis relied on *gyrB* phylotaxonomic marker sequences and their comparison to a reference database of all publicly available and taxonomically assigned sequences.

### Relationship between fertilization, soil origin, and rhizobium ZOTU richness and diversity in rooibos root nodules

Fertilization with sheep dung and soil origin had distinct effects on rhizobial richness and Simpson’s diversity only on *gyrB* communities (Table 2). Fertilized plants hosted a significantly higher number of different *gyrB* ZOTUs (47.81 ± 19.88 ZOTUs) than unfertilized plants (33.84 ± 16.99 ZOTUs) (Figure 3A; Table 2), which was also influenced, but to a lesser extent, by soil origin. Specifically, cultivated and mixed soils had significantly higher *gyrB* ZOTU richness compared to the uncultivated soils (Tukey’s HSD 95% confidence comparisons; Mixed-Cultivated: Estimate = 2.19 ± 2.77, P = 0.710; Mixed-Uncultivated: Estimate = 9.39 ± 2.75, P = 0.002; Cultivated-Uncultivated: Estimate = 7.20 ± 2.81, P = 0.030) (Table 2). Importantly, the effects of soil origin on root nodule community richness were independent from the fertilization treatment.

**Figure 3.**
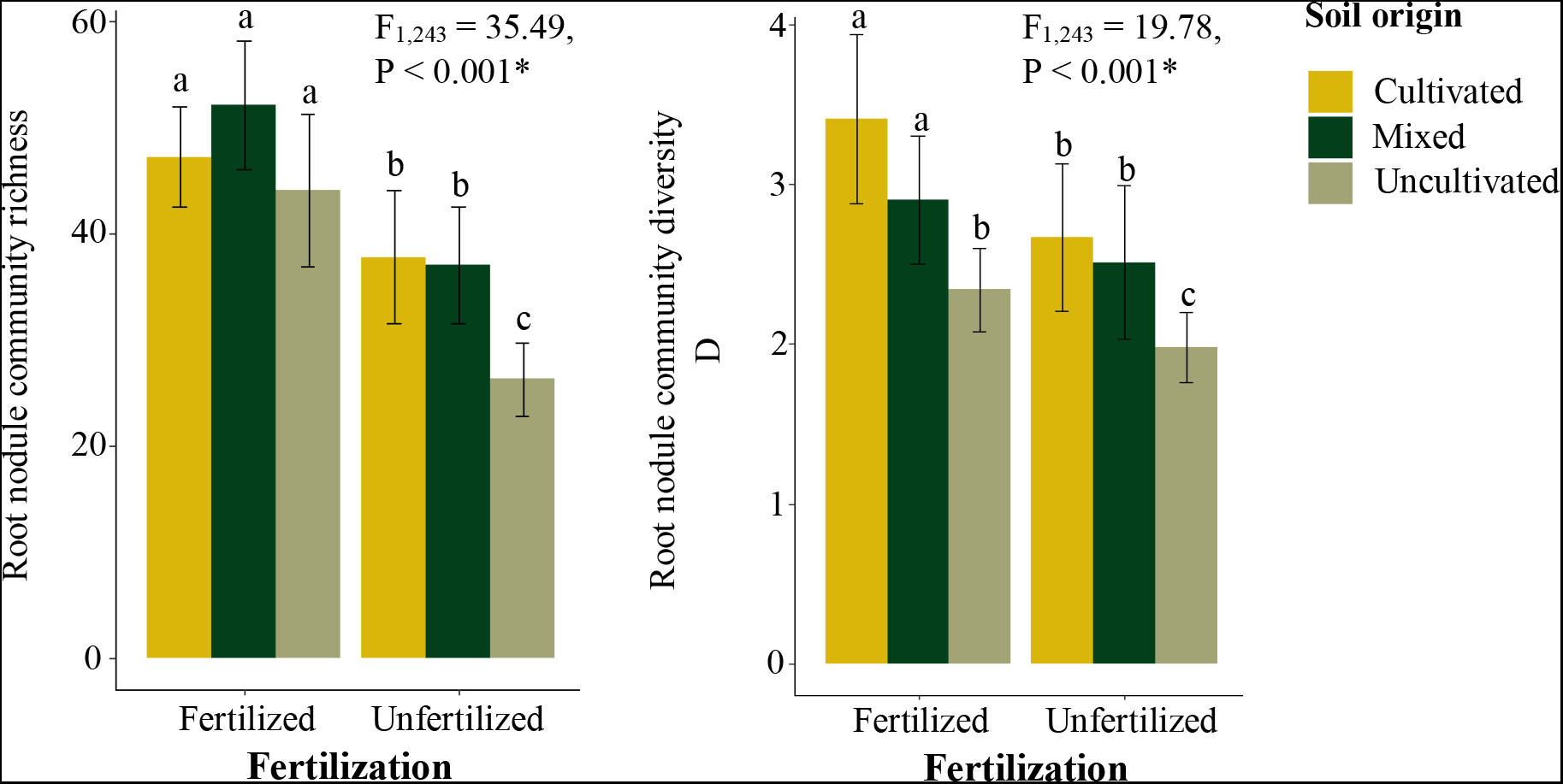
Effects of fertilization and soil origin on richness and diversity of bacterial operational taxonomic units (ZOTUs) in pools of nodules of rooibos grown in cultivated and wild soils and the mixture of these two soils. Soils were taken from five locations/farms, and were either fertilized with sheep dung or not. The ZOTUs were determined by similarity clustering of an 817 bp fragment of the phylotaxonomic marker gene *gyrB* at 99% sequence identity. The result of the mixed model testing the effects of fertilization on richness and diversity is reported on the graphs.

**Table 2.**
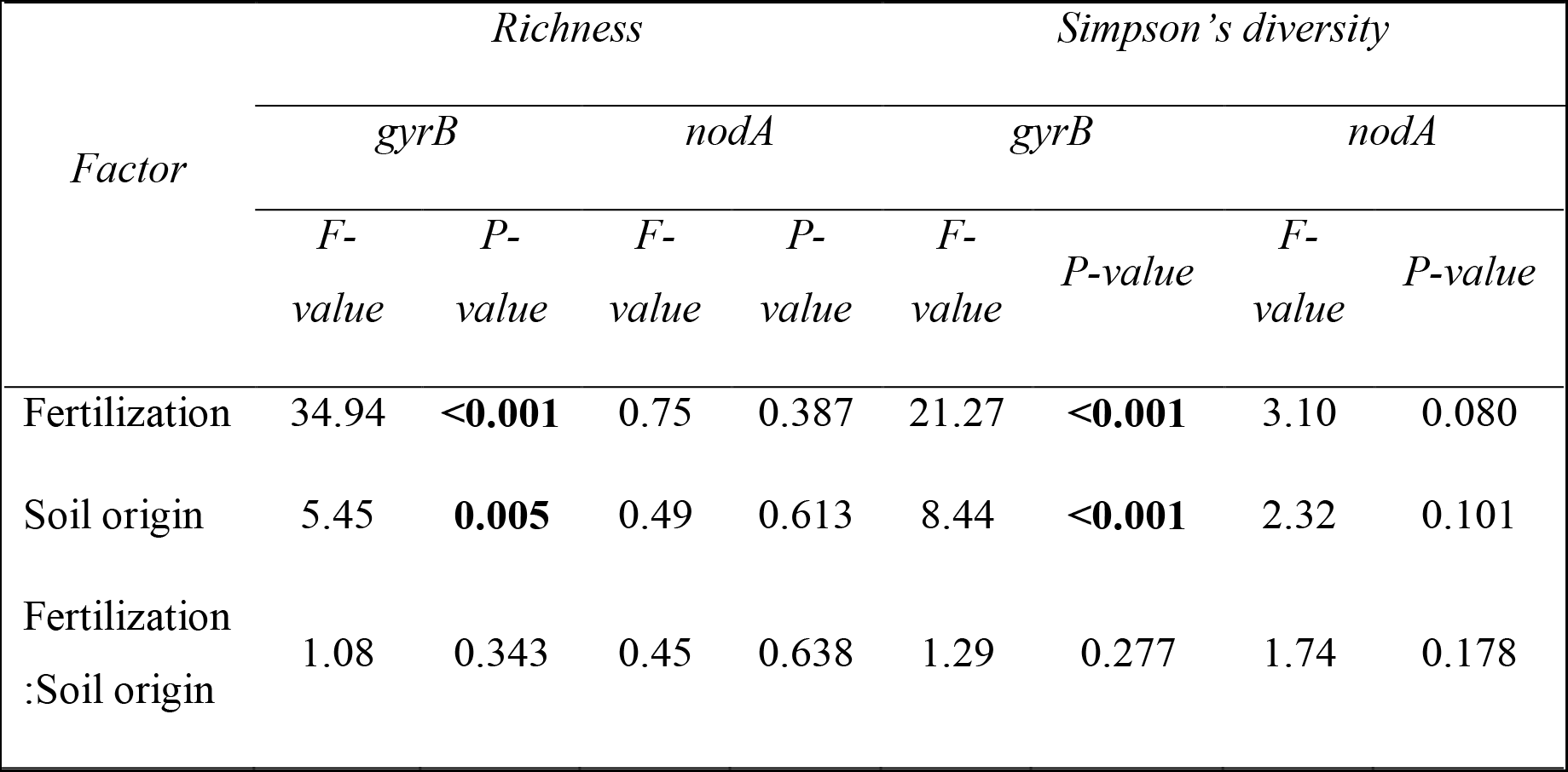
Results of mixed model analyses on the effects of fertilization, soil origin and their interaction on the richness and diversity of rhizobia in root nodules of rooibos. The root nodules were analysed using *gyrB* and *nodA* DNA sequences, which was used to calculate zero-sums operational taxonomic units (ZOTU) based on 99% sequence identity. Richness represents the total number of ZOTUs detected. The experimental factor ‘fertilization’ involved sheep dung addition to soil, and ‘soil origin’ comprises the use of soils from five pairs of plantations and adjacent uncultivated populations of rooibos and a 1:1 (v:v) mixture of those. The five locations (farms) were treated as a random and nested factor within fertilization and soil origin. There were 10 pot replicates per experimental treatment.

| *Factor* | *Richness* |  |  |  | *Simpson's diversity* |  |  |  |
| --- | --- | --- | --- | --- | --- | --- | --- | --- |
|  | *gyrB* |  | *nodA* |  | *gyrB* |  | *nodA* |  |
|  | *F-value* | *P-value* | *F-value* | *P-value* | *F-value* | *P-value* | *F-value* | *P-value* |
| Fertilization | 34.94 | **<0.001** | 0.75 | 0.387 | 21.27 | **<0.001** | 3.10 | 0.080 |
| Soil origin | 5.45 | **0.005** | 0.49 | 0.613 | 8.44 | **<0.001** | 2.32 | 0.101 |
| Fertilization<br>:Soil origin | 1.08 | 0.343 | 0.45 | 0.638 | 1.29 | 0.277 | 1.74 | 0.178 |

Plants grown on fertilized soils also had higher root nodule community diversity (D fertilized = 3.34 ± 2.23, D unfertilized = 2.39 ± 1.32) (Figure 3B; Table 2). Soil origin had a weak but significant effect on *gyrB* ZOTU community diversity as well. In this case, cultivated soils had significantly higher diversity than uncultivated ones, whereas mixed soils did not differentiate from the others (Tukey’s HSD 95% confidence comparisons; Mixed-Cultivated: Estimate = −0.536 ± 0.257, P = 0.095; Mixed-Uncultivated: Estimate = 0.554 ± 0.257, P = 0.082; Cultivated-Uncultivated: Estimate = 1.090 ± 0.262, P < 0.001) (Table 2). Like for community richness, the effects of soil origin on community diversity were independent from the fertilization treatment.

### Relationship between root biomass and the diversity of the root nodule-associated bacterial communities

The observed increase in *gyrB* ZOTU richness and diversity was positively correlated to root biomass (Figure 4A), as well as to the number of nodules per plant (a correlate of plant biomass) (Figure 4B; Supplementary Figure 1). The mixed effects model identified root biomass category as a significant factor explaining both root nodule community richness (F_2,191_= 12.54, P < 0.001) and diversity (F_2,193_= 5.05, P = 0.007). Post-hoc Tukey’s HSD tests confirmed larger root systems had significantly richer and more diverse root nodule communities than smaller ones (Supplementary Table 3).

**Figure 4.**
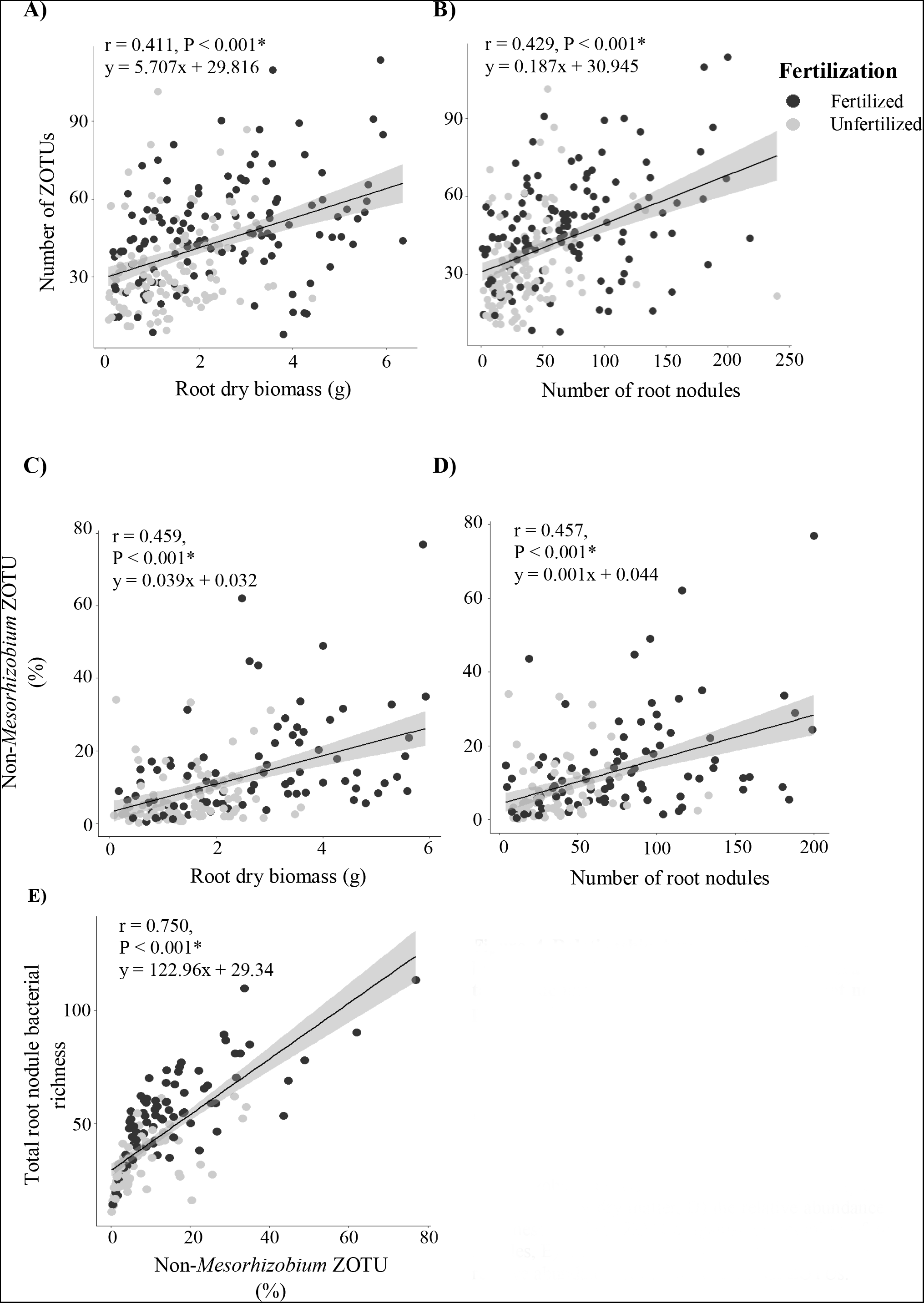
Relationship between the total number of bacterial (***gyrB***) and mesorhizobial (***nodA***) operational taxonomic units (ZOTUs) in single-plant root nodule pools and root biomass and the number of root nodules of eight-month-old rooibos plants. Seedlings were fertilized with sheep dung or not. Included are relationships between the contribution of non-mesorhizobial ZOTUs to total ZOTU diversity. Regression of A) the number of ZOTUs against the root system size, B) the number of ZOTUs against the number of root nodules, C) the relative abundance of non-mesorhizobial ZOTUs extracted from the gyrB sequences against the root dry matter, D) the relative abundance of non-mesorhizobial ZOTUs against the number of root nodules, E) the total bacterial ZOTU richness against the relative abundance of non-mesorhizobial ZOTUs.

Since increased Simpson’s diversity implied reduced dominance by *Mesorhizobium*, we correlated the abundance of non-*Mesorhizobium* ZOTUs to root biomass and the number of root nodules per plant. Non-*Mesorhizobium* ZOTUs became more abundant with larger root systems and higher numbers of nodules (Figure 4 C, D). When the relative abundance of non-*Mesorhizobium* strains was directly correlated to richness, the correlation coefficients became even larger (Figure E). This pointed at a prominent contribution of rare ZOTUs to the overall alpha diversity and root nodule community structure of rooibos. Importantly, the strong correlation between non-*Mesorhizobium* relative abundance and the overall root nodule community diversity indicates such rare ZOTUs reduce the dominance of *Mesorhizobium* in larger root systems. We then explored the relative contribution of such non-*Mesorhizobium* ZOTUs to the observed richness increase with larger root systems compared to the dominant *Mesorhizobium*. We found a much stronger correlation between non-*Mesorhizobium* ZOTU richness and the overall root nodule community richness (r = 0.945, P < 0.001), compared to *Mesorhizobium* richness (r = 0.381, P < 0.001).

The genera that contributed the most to the higher root nodule *gyrB* community richness in larger root systems were identified using indicator species analysis. In descending order, the top indicator ZOTU genera from plants with large root systems were *Bradyrhizobium* (20 ZOTUs), *Rhizobium* (12 ZOTUs), and *Herbaspirillum* (8 ZOTUs) (all selected based on correlation coefficients r > 0.8 and P < 0.05). Noteworthy, there were no indicators from medium-sized root systems, while 6 ZOTUs were associated with small root systems. From these 6 ZOTUs, the only 2 ZOTUs that were identified at genus level were both *Bradyrhizobium*, and were only found in a few samples from two particular locations.

Since most of the plants with larger root systems that produced more nodules were from the fertilized treatment, we assessed whether the increase in non-*Mesorhizobium* ZOTU relative abundance could be due to additional taxa introduced directly with the fertilizer (i.e. fertilizer-borne taxa). This confirmed that environmental preference for nutrient-rich soil conditions was an unlikely explanation for the increased richness observed in larger root systems. This is so because, from the 10 ZOTUs that were unique to the fertilized treatment, the one with highest (yet very low) cumulative relative abundance belonged to the genus *Mesorhizobium* (17.56% in cumulative relative abundance), the others being 5.37% or less (Supplementary Table 4). Moreover, these fertilizer-related ZOTUs occurred in very few samples, indicating their low occurrences in *gyrB* communities within fertilized treatments (Supplementary Figure 2). The fact that only 10 ZOTUs were unique to fertilized plants and their rarity suggests a limited role of environmental preferences in explaining the increased rhizobium diversity in larger root systems.

### Beta diversity patterns of root nodule communities in rooibos

NMDS ordination plots based on Bray-Curtis *nodA* dissimilarities revealed strong community structure associated with the location from which soils were collected (Figure 5 A-C). Geographical location seemed less important in driving community structure derived from *gyrB* compared to *nodA* (Figure 5 D-F). Mixing of soils from the same location led to an even stronger geographical signal in community dissimilarity (Figure 5C). The PERMANOVA results confirmed the geography-driven β diversity patterns from the ordination plots, also showing that soil origin influenced community composition for both *gyrB* and *nodA* datasets (Table 3). Based on the R^2^ values, soil origin was twice as important in structuring *gyrB* than *nodA* communities. Moreover, fertilization had a weak but significant effect on *gyrB* community structure, but no effect on *nodA* communities. Small effects on β diversity were also found for root system size, while there were no effects from the interaction between fertilization and soil origin (Table 3). Despite high stress values, the NMDS plots provide informative insights given the agreement with the PERMANOVA results.

**Figure 5.**
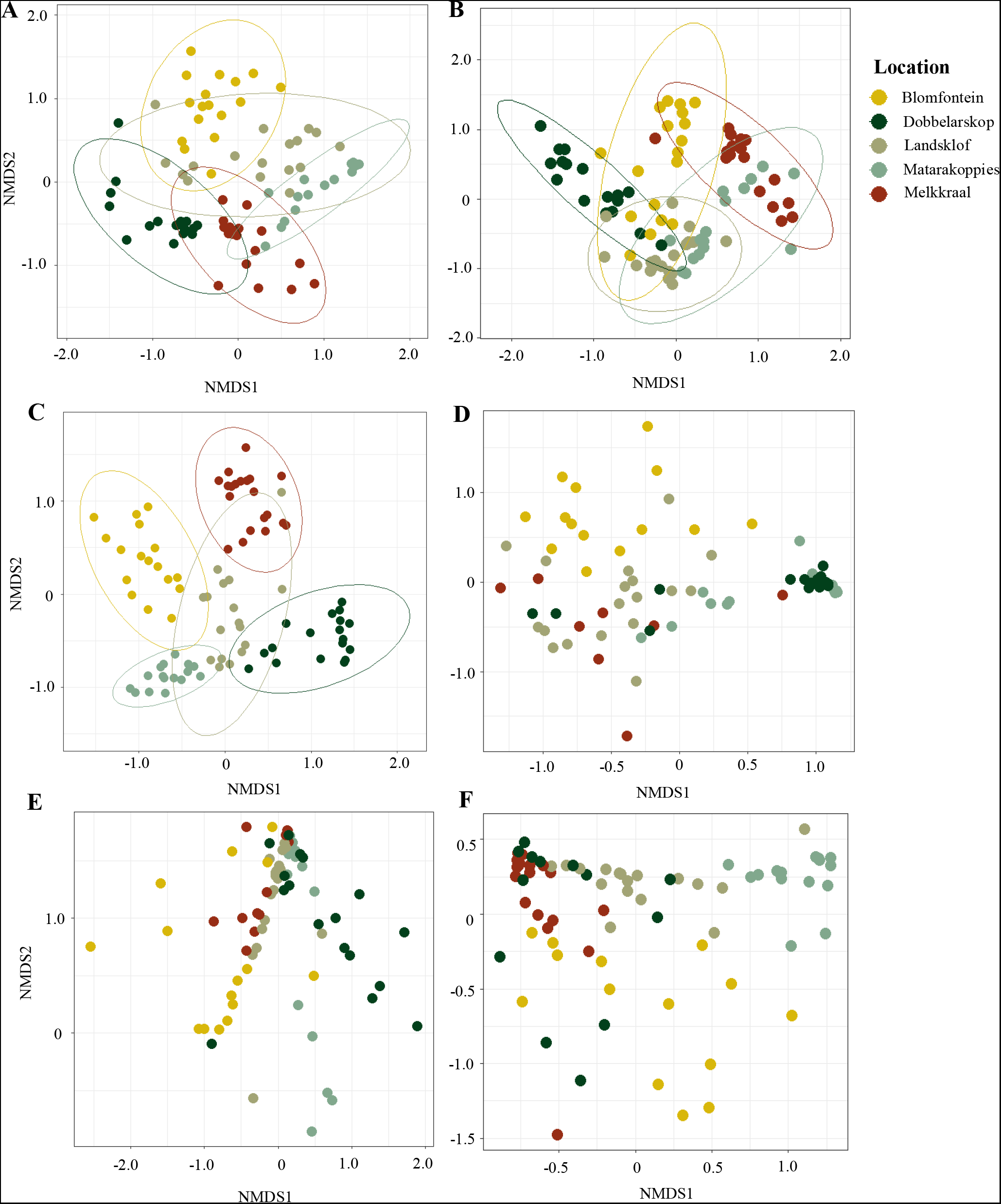
Non-metric multidimensional scaling (NMDS) of Bray-Curtis dissimilarities between the root nodule community structures from pools of root nodules of single rooibos plants grown in soil from five locations/farms for eight months. Half of the plants were fertilized with sheep dung. Root nodule community structures were characterized using the nodulation gene, *nodA*, and the DNA gyrase subunit B phylotaxonomic marker, *gyrB*. Ordination plot of bacterial communities analysed with *nodA* from cultivated (pane A, Stress = 0.24) and uncultivated soils (pane B, Stress = 0.23). C) Ordination plot of bacterial communities of plants grown in a 1:1 (v:v) mixture of soil from plantations and adjacent uncultivated populations of rooibos analysed with *nodA* (Stress = 0.22). Ordination plot of root nodule community structures analysed with *gyrB* from cultivated (pane D, Stress = 0.16) and uncultivated soils (E, Stress = 0.16). F) Ordination plot of bacterial communities of plants grown in a 1:1 (v:v) mixture of soil from plantations and adjacent uncultivated populations of rooibos analysed with *gyrB* (Stress = 0.14), 95% confidence ellipses are shown when differences between clusters explained more than 10% of the variation.

**Table 3.** Results of a permutational analysis of variance (PERMANOVA) on the effects of location, soil origin, fertilization and root system size and their interactions on the rhizobial community structures in root nodules of rooibos). DNA sequence data of the *gyrB* and *nodA* genes were used to determine operational taxonomic units at 99% sequence identity whose relative abundances were used to calculate Bray-Curtis dissimilarities. The analysis was run with 9999 permutations.

| *Factor* | *nodA* |  |  |  | *gyrB* |  |  |
| --- | --- | --- | --- | --- | --- | --- | --- |
|  | *df* | *F-value* | *R*<sup>2</sup> | *P-value* | *F-value* | *R*<sup>2</sup> | *P-value* |
| Location | 4 | 30.28 | 0.246 | **<0.001** | 17.56 | 0.174 | **<0.001** |
| Soil origin | 2 | 10.29 | 0.042 | **<0.001** | 14.29 | 0.071 | **<0.001** |
| Fertilization | 2 | 1.27 | 0.003 | 0.204 | 3.36 | 0.008 | **0.010** |
| Root system size | 2 | 2.01 | 0.008 | **0.003** | 2.64 | 0.013 | **0.008** |
| Location:Soil origin | 8 | 11.88 | 0.193 | **<0.001** | 10.19 | 0.202 | **<0.001** |
| Fertilization:Soil<br>origin | 2 | 1.28 | 0.004 | 0.171 | 1.58 | 0.008 | 0.106 |

In order to explain the observed β diversity differences, we calculated the distances between centroids of the location × soil clusters based on Bray-Curtis distances for both genes and correlated them to geographical distances between locations and the euclidean distances between soil properties. We found a significant positive correlation (r = 0.515, P < 0.001) between geographical distances and centroid distances for the *nodA* marker (Figure 6A), but not for the *gyrB* communities (r = 0.175, P = 0.626; Figure 6B). For the *nodA* communities we only found a significantly positive correlation of distances between centroids and soil Mg concentration (r=0.227, P=0.02). For *gyrB*, significant correlations were found for water holding capacity (r=0.303, P=0.002; Figure 6C, D), soil K concentration (r=0.282, P=0.004), and soil N concentration (r=0.200, P=0.045).

**Figure 6.**
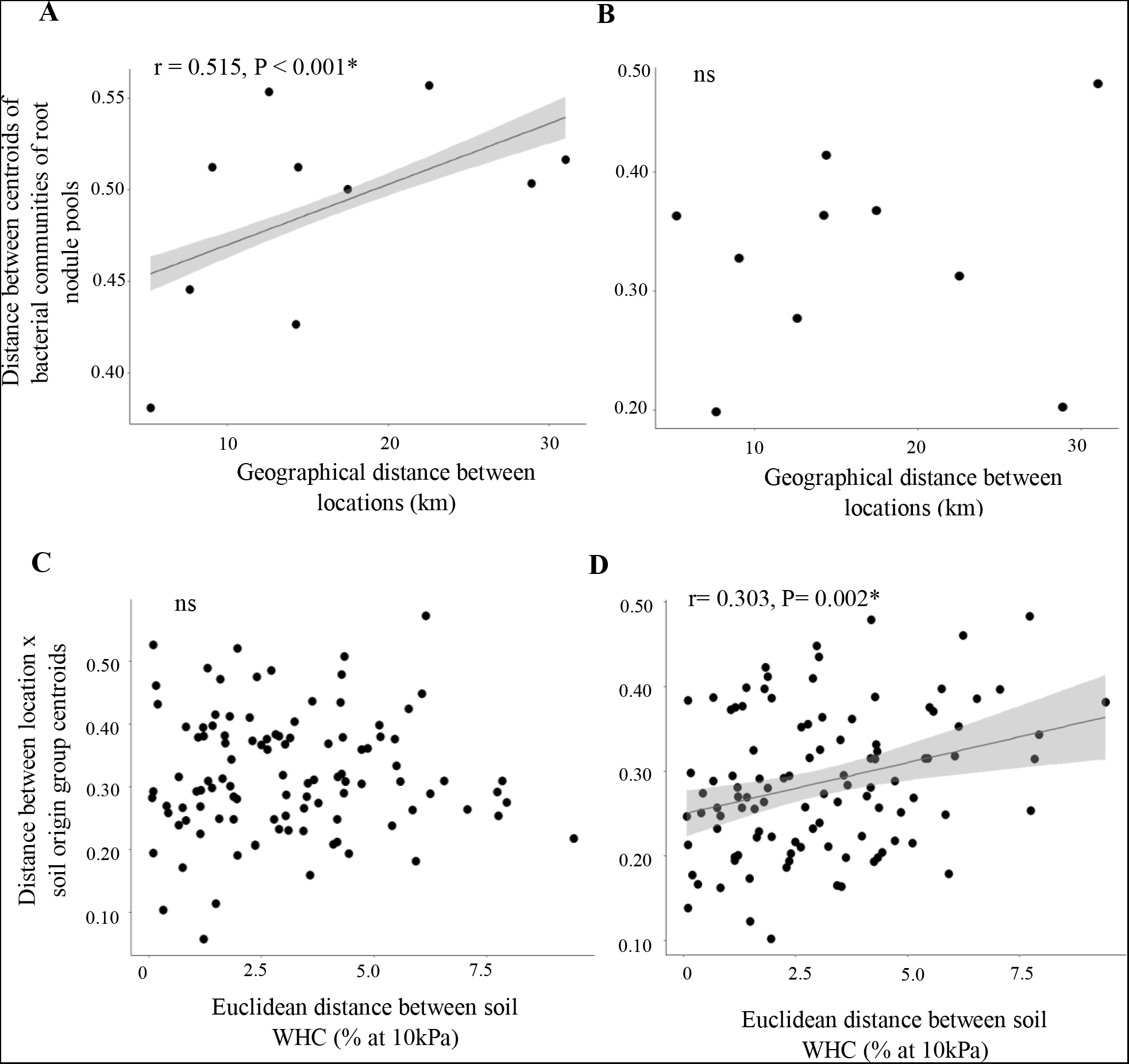
Relationship between the differences among the rhizobial community structures of root nodule pools of rooibos and geographical distances between locations (A, B), and water holding capacities (WHC) of different soils (C, D). The rhizobial community structures were characterized using the nodulation gene nodA (A, C) and the DNA gyrase subunit B phylotaxonomic gene, gyrB (B, D).

Finally, following the methods of Stegen et al., (2015), we calculated the relative contribution of neutral (drift and dispersal-based) and deterministic (selection-based) processes in driving species turnover in core symbiotic and bacterial root nodule communities of rooibos. We found a predominance of neutral processes in driving both *gyrB* and *nodA* community turnover (Fig. 7). Root nodule bacterial communities (*gyrB*) were driven mostly by ecological drift (62.6%), followed by homogeneous selection (25.6%), and the combination of dispersal limitation and drift (6.4%). In contrast, the core symbiotic communities of rooibos (*nodA*) were mostly affected by dispersal limitation and drift (82.7%), followed by drift alone (15.1%) (Fig. 7).

**Figure 7.**
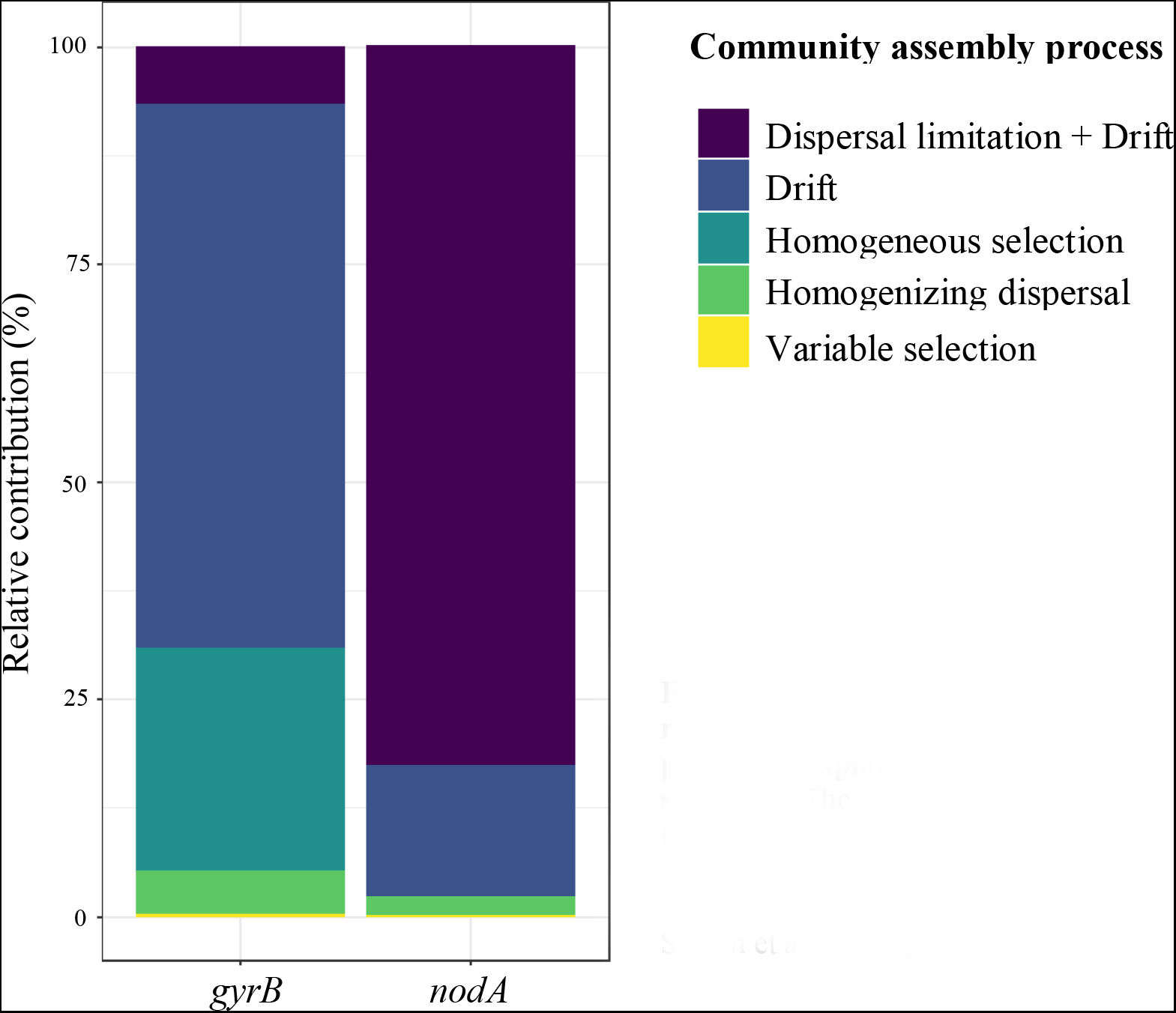
Relative contribution of neutral (drift and dispersal-related) and deterministic (selection-related) ecological processes explaining differences in root nodule community structure. The core rhizobial community was characterized using the nodulation gene nodA and the bacterial root nodule community was characterized using the gyrB gene. Proportions are based on β-NTI and RC_Bray_ values obtained by comparison to null models as in Stegen et al., (2015).

## Discussion

In this study we asked whether bacterial communities assemble in neutral or deterministic ways in the root nodules of a promiscuous legume crop adapted to extremely low-nutrient soils. While in rooibos both deterministic and stochastic processes contribute to root nodule community assembly, we found evidence to suggest that it is mostly neutral. Importantly, the core rhizobial symbionts of rooibos (*Mesorhizobium*) were most affected by dispersal limitation in combination with drift, whereas other bacterial members of the root nodule community were more affected by drift alone and selection. Distinct changes in taxonomic diversity with root system biomass, and strong geographical structure in the *Mesorhizobium* group, support such findings.

Contrary to what has been found in most legumes, fertilized rooibos plants produced more nodules on average, and importantly, these harboured richer and more diverse bacterial communities. We suggest that nodule production that is proportional to root biomass production is unsurprising given the low nutrient availability typical of Fynbos soils. This can be seen as an ecological insurance (or “bet-hedging”) strategy (Yachi and Loreau, 1999; Starrfelt and Kokko, 2012), in which initial resource allocation into more nodules may ensure some symbiotic benefits, whether optimal or not, under future conditions of extreme nutrient limitation. Physiological and phenotypic strategies of plants closely related to rooibos with similar adaptations to post-fire recruitment, suggest that such bet-hedging strategies may be common under nutrient-poor conditions (e.g. within the Podalyrieae tribe: *Cyclopia spp.* and *Podalyria spp.*; Maistry *et al.*, 2013; Mndzebele and Dakora, 2017). Such strategy may be related to root nodule bacterial diversity as well (see the balancing nodulation hypothesis, Siler and Friesen, 2017).

Root system biomass strongly correlated with the diversity of bacterial taxa that are not the core symbionts of rooibos. This positive trend happened independently of the soil abiotic conditions, implying it cannot be due to certain strains having environmental preference for particular conditions, but rather a direct consequence of larger root systems or a better health status of the plant (Dinnage *et al.*, 2019). Mostly rhizobial taxa (*Rhizobium, Bradyrhizobium, Agrobacterium*), and groups known for their nitrogen-fixation ability (*Herbaspirillum*), colonized nodules from larger root systems. We interpret this as a commensal interaction, whereby taxa supported by neighbouring vegetation may take advantage of the poor ability of rooibos to control nodule infection (Pahua *et al.*, 2018). Under the ecological insurance hypothesis, this *a priori* costly interaction may reward the plant in the form of improved nutrition from drought- or nutrient limitation-adapted taxa on the long run.

Similar to our findings, Dinnage *et al.*, (2019) recently reported increased root nodule rhizobial richness to be linked with root system size in the Australian legume *Acacia acuminata*. This study, based on field data from plants of different age classes, pointed at a role of niche construction during plant growth that would favour the accumulation of rhizobial diversity over time. Under this assumption, root development would make new resources available that could be used by an increasingly wider range of rhizobium strains. While plausible in *Acacia acuminata*, this option is less likely to apply to the present study, as harvested rooibos seedlings were evenly aged and grown in pots. However, Dinnage *et al.*, (2019) also pointed to the role that neutral processes may play in driving the establishment of richer rhizobial communities in larger root systems, which would operate as “islands” (i.e. Theory of Island Biogeography, McArthur and Wilson, 2001). Within such framework, the probability of establishment and maintenance of root nodule bacterial taxa would be proportional to the size of the root system (Dinnage *et al.*, 2019), underlying a neutral community assembly. Support for this island biogeographic model of microbial community assembly is strong from both the ◻- and ◻- diversity perspective of the observations made in our study, as discussed below.

The communities of *Mesorhizobium* and rare bacterial taxa in the root nodules were mostly structured by the neutral process of drift, but operating at different spatial scales. The *nodA* communities (true symbionts of rooibos) were mostly structured by geographical origin, which agrees with the predominance of dispersal limitation together with drift in explaining species turnover in these communities. This is further supported by the positive distance-decay in community dissimilarity in the *nodA* community, which is a known indicator of ecological drift (Hanson *et al.*, 2012). This suggests that *nodA* assemblages are most likely shaped by dispersal limitation between distinct rooibos populations. In fact, as a functional marker, the evolution of *nodA* is closely linked to that of its associated host plants (Masson-Boivin *et al.*, 2009; Lemaire *et al.*, 2015). Given the habitat heterogeneity characteristic of the Core Cape Subregion (Cowling *et al.*, 2008), and the known diversity of rooibos ecotypes (Malgas *et al.*, 2010; Hawkins *et al.*, 2011), drift is likely to drive rhizobial community differences between partially isolated rooibos populations (Sprent *et al.*, 2017).

In contrast, the *gyrB* communities, which include a myriad of non-symbiotic taxa, were almost entirely driven by drift alone. When not combined with dispersal limitation, drift underlies the absence of environmental filters (Dumbrell *et al.*, 2010), in this case the absence of filtering by the plant host. As shown before, non-*Mesorhizobium* taxa accumulated proportionally to root system biomass, which agrees with the dominance of drift as there were no environmental filters to root nodule colonization. A proportion of species turnover in these communities was driven by homogenizing selection. Here again, this agrees with the positive relationships observed between *gyrB* community dissimilarity and soil abiotic factors such as WHC. It is known that environmental filtering can favour strains with distinct environmental preferences, for example in response to soil N and P (Vuong *et al.*, 2017), cropping practices (Yan *et al.*, 2014), or surrounding vegetation (Slabbert *et al.*, 2010; Pahua *et al.*, 2018).

We did not find evidence that differences in rhizobium root nodule communities affect the performance of rooibos plants (Figure S3). The maintenance of *Mesorhizobium* dominance throughout the treatments suggests that rooibos derives most of its symbiotic benefits from this genus. Experimental evidence shows that inoculating plants with diverse rhizobial strains decreases symbiotic benefit due to rhizobial strain competition over the short term (Simonsen and Stinchcombe, 2014; Barrett *et al.*, 2015). Despite the action of such biotic interactions within nodules, legumes favour the establishment and replication of the best performing strains (Kiers *et al.*, 2003). When contrasted with the evidence for the “balancing nodulation” hypothesis (Siler and Friesen, 2017), it seems that legumes favour the dominance of very few effective strains to avoid strain competition, while a wide diversity is maintained at very low densities as a potential insurance for the case that alternative symbionts become more beneficial under changing conditions. This complies with the idea that, over the long run, rooibos favours the fitness of *Mesorhizobium* strains within the root nodules, despite a large diversity of other taxa is neutrally maintained at densities low enough to avoid strain competition (Le Roux *et al.*, 2017). Such prediction would be confirmed by studying root nodule assemblages of rooibos in the transition to the nutrient-poor conditions characteristic of its mature life stages, or by performing field studies on distinct mature populations.

## Conclusions

Root nodule community assembly in rooibos seedlings is dominated by ecological drift, but it operates at different spatial scales for the core rhizobial symbionts and apparently opportunistic bacterial taxa. Specifically, at the root level, an accumulation of rare bacterial taxa dominated by ecological drift occurs proportionally to the biomass of the root system. Instead, the core rhizobial symbionts of rooibos are structured by dispersal limitation and drift, which operate on a regional scale. The “balancing nodulation” hypothesis (Siler and Friesen, 2017), purports that legumes obtain some long-term benefit from associating with high rhizobium diversity at low densities, such as flexibility to shift the association from one to another symbiont under changing environmental conditions. Our findings suggest under low symbiont selectivity this diversity is maintained by the neutral process of drift. By relating ecological pattern to process, this study shows that dominant symbiotic and rare rhizobial taxa can be structured by different ecological processes. Accounting for these processes can contribute to maintaining root symbiont diversity under productive agricultural conditions, despite the role of root nodule diversity for plant functioning needs to be clarified in the future.

## Acknowledgements

This study was funded by the Mercator Research Program of the World Food Systems Center of ETH Zurich. Data produced and analyzed in this paper were generated in collaboration with the Genetic Diversity Center of ETH Zurich (GDC) and the Functional Genomics Center of the University of Zurich (FGCZ). JR acknowledges the assistance from Dr. Jean-Claude Walser, Dr. Andrea Patrignani and Dr. Weihong Qi. JLR acknowledges funding from Macquarie University’s Faculty of Science and Engineering and Department of Biological Sciences. Finally, the authors acknowledge the assistance of Noel Oettlé, Dr. Cecilia Bester, and the community of rooibos farmers in the Suid Bokkeveld for making the sampling in South Africa possible.

