## Supplementary Material for "Different ecological processes drive the assembly of dominant and rare root-associated bacteria in a promiscuous legume"

**Figure S1 Effect of fertilization on A) the number of root nodules and B) the relationship between the root nodule number and root dry biomass.** Boxplots show the median and first and third quartile and the whiskers the range of 1.5 standard deviations. The correlation coefficient and significance of the linear regression are indicated. For the statistical results of Suppl. Fig. 1A, see Table 1.

**A**

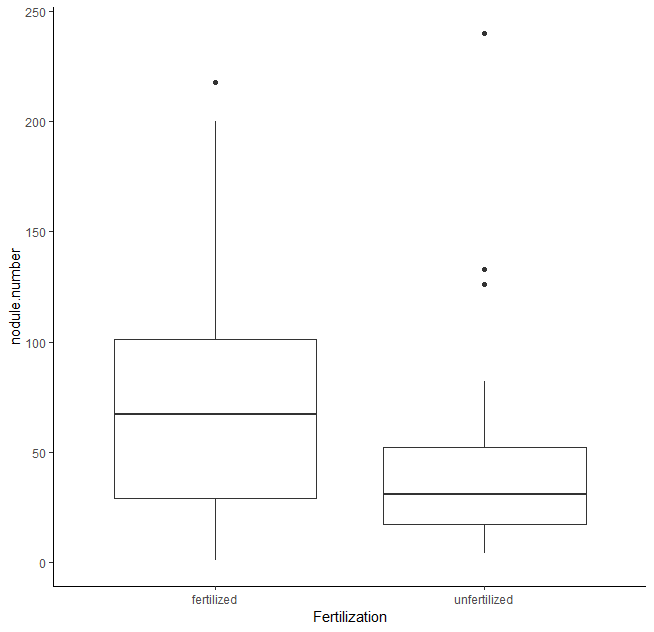

**Fertilization**

Fertilized

Unfertilized

0

50

100

150

200

250

Number of root nodules

F_1,271_ = 63.57, P < 0.001

**B**

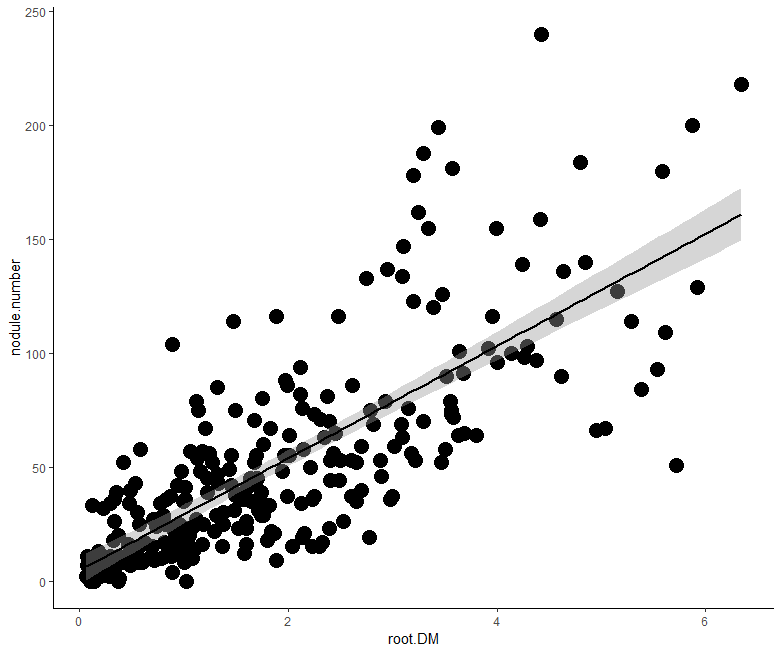

0

2

4

6

Root biomass (g)

0

50

Number of root nodules

100

150

200

250

r = 0.781, P < 0.001

**Figure S2 Occurrence and relative abundance of indicator operational taxonomic units for fertilized rooibos plants. ZOTUs were considered indicators when the correlation coefficient (r) between their relative abundance and frequency in a particular treatment was more than 0.8 and at statistical significance of P < 0.05.** The taxonomic affiliations are shown at the left and the relative abundances are indicated in blue tones. n = 116.

*Herbaspirillum*

*Rhizobium*

*Unidentified Rhizobiaceae*

*Mesorhizobium*

*Brevundimonas*

*Bradyrhizobium*

*Unidentified Alpha- proteobacterium*

Samples (n=116)

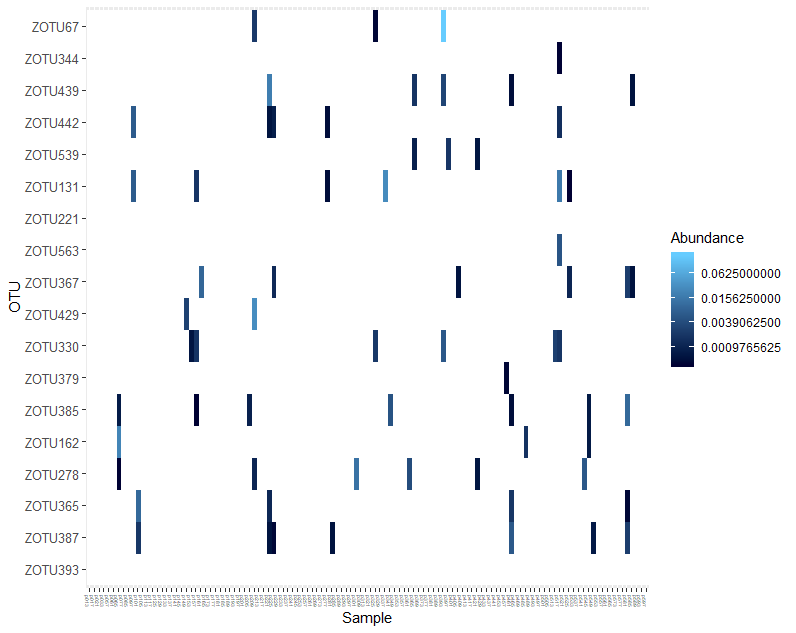

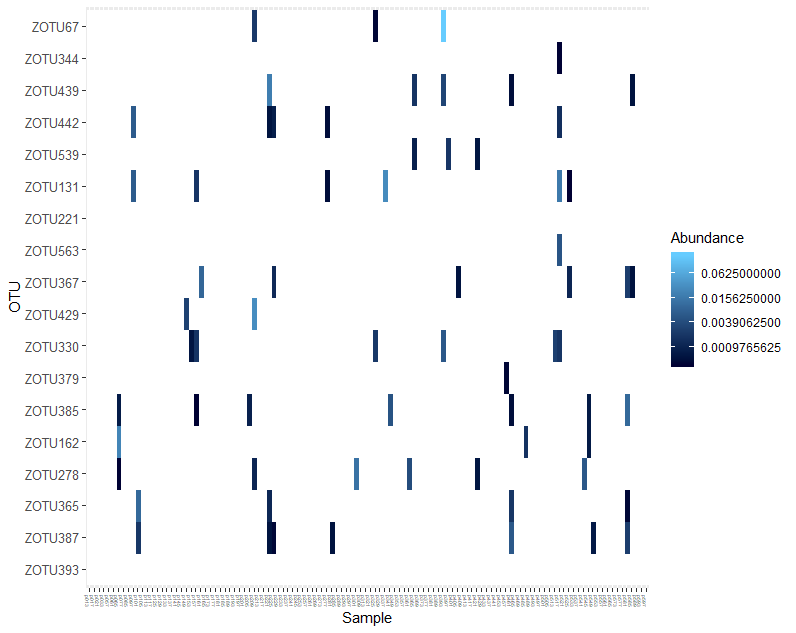

Relative abundance (%)

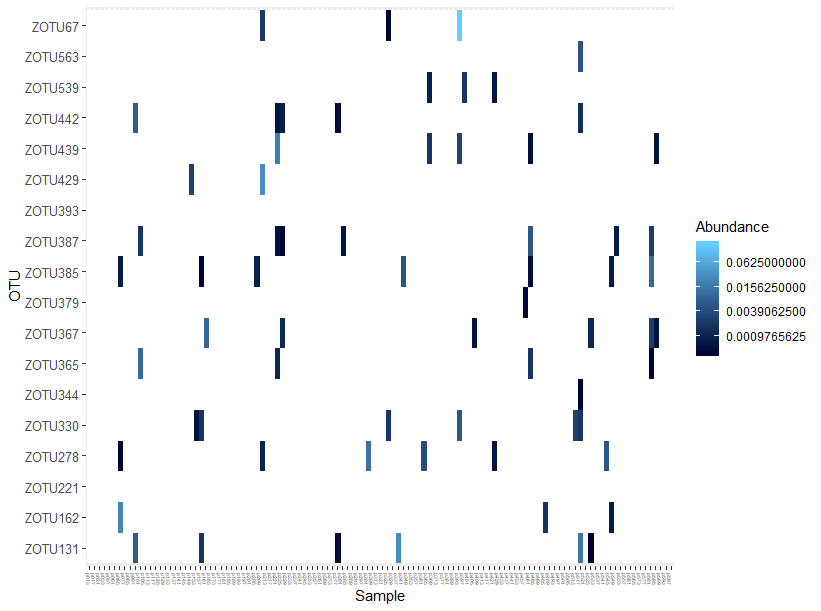

17.5

0

Cultivated

Mixed

Uncultivated

**Figure S3 Canonical analysis of principal coordinates (CAP) of the rhizobial community structures in pools of root nodules of rooibos and plant growth physiological parameters.** Bray-Curtis distances of the bacterial communities and total plant dry matter, nodule number, nodule density (i.e. ratio between nodule number and root dry biomass), leaf nitrogen (N) to phosphorus (P) ratio, and foliar δ^15^N signatures are displayed. The root nodule community structures were characterized using the nodulation gene, *nodA* (A) and the DNA gyrase subunit B phylotaxonomic gene, *gyrB* (B).

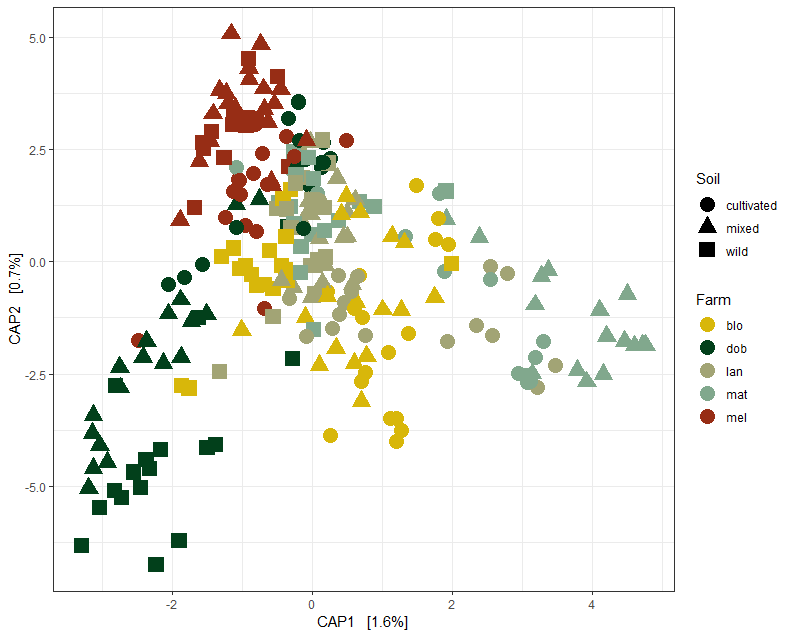

CAP2 (0.7%)

0

2.5

5.0

-5.0

-2.5

CAP1 (1.6%)

0

2.0

4.0

-2.0

Plant biomass

Nodule number

Foliar ∆15N

Foliar N:P ratio

Nodulation

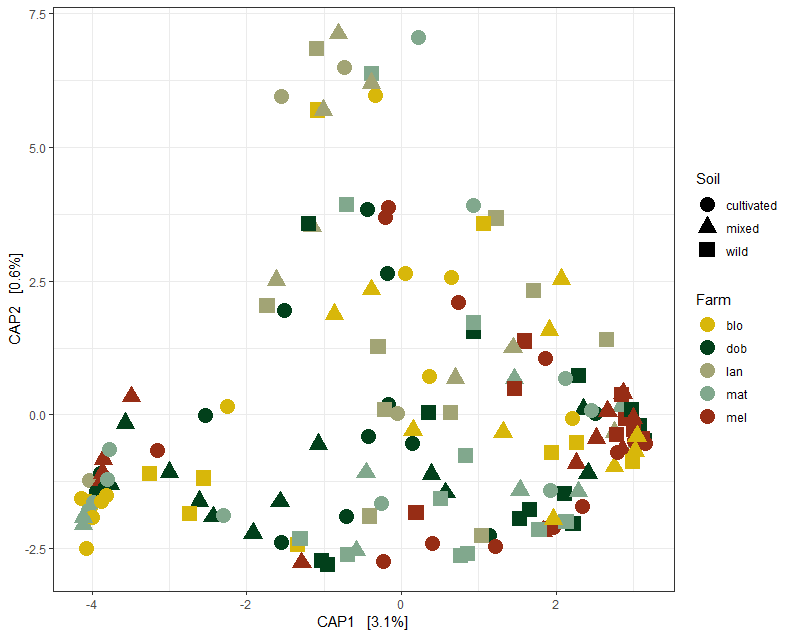

Foliar N:P ratio

Nodulation

Nodule number

Plant biomass

Foliar ∆15N

CAP2 (0.6%)

5.0

7.5

2.5

0

-2.5

CAP1 (3.1%)

-4.0

-2.0

2.05

0

Blomfontein

Dobbelarskop

Landsklof

Matarakoppies

Melkkraal

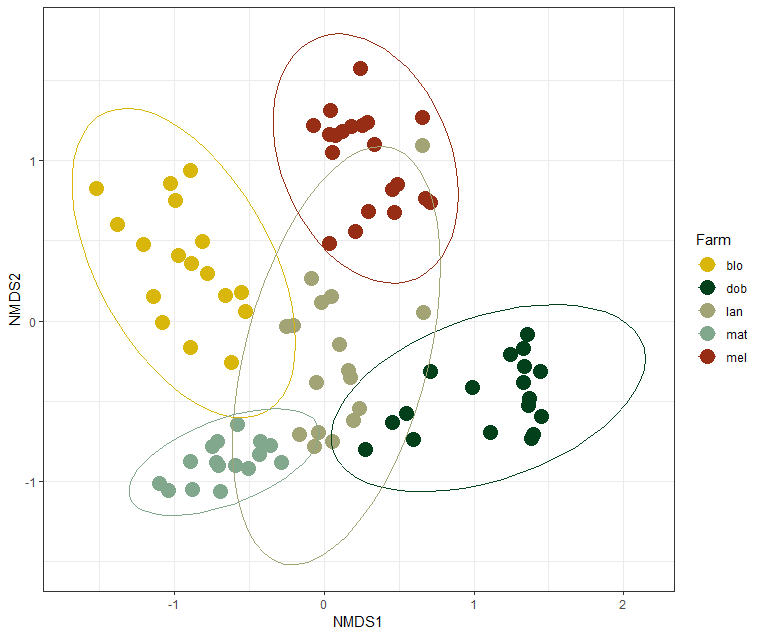

**Location**

Mixed

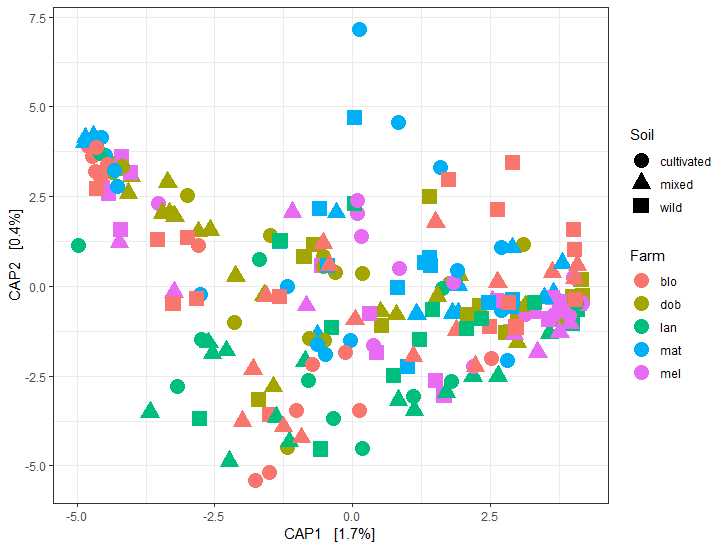

**Soil**

Cultivated

Uncultivated

**Table S1 Physicochemical properties of soils from five plantations and adjacent uncultivated populations of rooibos collected from cultivated and uncultivated rooibos populations and sheep dung fertilizer.** Locations/farms: Blo, Blomfontein; Dob, Dobbelarskop; Lan, Landsklof; Mat, Matarakoppies; Mel, Melkkraal.

| *Soil properties* | *Fertilizer (sheep dung)* | *Cultivated soil (plantations)*  *(Locations/farms)* | | | | | *Uncultivated soil (wild populations)*  *(Locations/farms)* | | | | |
| --- | --- | --- | --- | --- | --- | --- | --- | --- | --- | --- | --- |
|  |  | *Blo* | *Dob* | *Lan* | *Mat* | *Mel* | *Blo* | *Dob* | *Lan* | *Mat* | *Mel* |
| CEC (cmol kg^-1^) | - | 1.42 | 1.30 | 1.54 | 1.43 | 1.76 | 1.75 | 1.85 | 1.28 | 1.60 | 1.42 |
| WHC (% at 10 kPa) | - | 13.79 | 15.76 | 12.09 | 17.23 | 20.04 | 17.55 | 21.53 | 16.81 | 15.60 | 19.85 |
| Clay (%) | - | 7 | 5 | 5 | 5 | 9 | 7 | 7 | 5 | 7 | 7 |
| pH (CaCl_2_) | - | 5.78 | 5.76 | 6.15 | 5.53 | 5.55 | 5.89 | 5.48 | 5.51 | 5.60 | 5.29 |
| Organic C (%) | - | 0.54 | 0.65 | 0.54 | 0.75 | 1.11 | 1.21 | 1.79 | 0.87 | 3.93 | 1.05 |
| N (g kg^-1^) | 15.93 | 0.11 | 0.13 | 0.11 | 0.14 | 0.22 | 0.26 | 0.37 | 0.15 | 0.16 | 0.22 |
| P (g kg^-1^) | 1.86 | 0.020 | 0.001 | 0.003 | 0.010 | 0.007 | 0.005 | 0.030 | 0.007 | 0.010 | 0.020 |
| K (g kg^-1^) | 8.70 | 0.38 | 0.53 | 0.24 | 0.54 | 1.22 | 0.57 | 1.22 | 1.01 | 0.82 | 0.81 |
| ^15^N (‰) | 16.36 | 20.40 | 19.75 | 23.68 | 22.29 | 14.52 | 10.87 | 10.31 | 20.26 | 19.31 | 13.91 |
| N:P ratio | 9.53 | 4.98 | 10.94 | 36.17 | 13.91 | 33.06 | 47.84 | 12.74 | 20.80 | 16.07 | 10.91 |
| Ca (g kg^-1^) | 12.30 | 0.06 | 0.11 | 0.06 | 0.05 | 0.12 | 0.14 | 0.16 | 0.06 | 0.06 | 0.08 |
| Mg (g kg^-1^) | 7.55 | 0.10 | 0.17 | 0.12 | 0.07 | 0.29 | 0.17 | 0.35 | 0.12 | 0.07 | 0.15 |

**Table S2 Results of mixed model analyses on the effects of fertilization, soil origin and their interaction on leaf and root macro- and micronutrient contents of rooibos.** The experimental factor ‘fertilization’ involved sheep dung addition to soil, and ‘soil origin’ comprises the use of soils from five pairs of plantations and adjacent uncultivated populations of rooibos and a 1:1 (v:v) mixture of those. The five locations (farms) were treated as a random and nested factor within fertilization and soil origin. There were 10 pot replicates per experimental treatment, n=271.

| *Response variable* | *Experimental factors* | | | | | |
| --- | --- | --- | --- | --- | --- | --- |
|  | *Fertilization (df = 2)* | | *Soil origin (df = 2)* | | *Fertilization:Soil origin (df = 2)* | |
|  | *F-value* | *P-value* | *F-value* | *P-value* | *F-value* | *P-value* |
| Leaf N (mg g^-1^) | 8.62 | **0.004** | 0.01 | 0.998 | 2.36 | 0.100 |
| Leaf P (mg g^-1^) | 102.05 | **<0.001** | 6.30 | **0.002** | 0.01 | 0.994 |
| Leaf K (mg g^-1^) | 7.38 | **0.007** | 5.87 | **0.003** | 3.60 | **0.029** |
| Leaf Ca (mg g^-1^) | 0.42 | 0.516 | 6.72 | **0.002** | 1.57 | 0.210 |
| Leaf Mg (mg g^-1^) | 6.47 | **0.012** | 13.10 | **<0.001** | 1.35 | 0.261 |
| Leaf Mn (mg g^-1^) | 17.20 | **<0.001** | 0.14 | 0.865 | 1.61 | 0.202 |
| Leaf Fe (mg g^-1^) | 10.50 | **0.001** | 1.01 | 0.366 | 1.59 | 0.206 |
| Root N (mg g^-1^) | 51.42 | **<0.001** | 1.23 | 0.294 | 0.64 | 0.526 |
| Root P (mg g^-1^) | 159.77 | **<0.001** | 3.54 | **0.031** | 0.19 | 0.829 |
| Root K (mg g^-1^) | 257.47 | **<0.001** | 0.147 | 0.864 | 0.33 | 0.720 |
| Root Ca (mg g^-1^) | 93.77 | **<0.001** | 1.18 | 0.309 | 2.37 | 0.100 |
| Root Mg (mg g^-1^) | 84.86 | **<0.001** | 1.89 | 0.154 | 0.85 | 0.429 |
| Root Mn (mg g^-1^) | 55.25 | **<0.001** | 9.61 | **<0.001** | 4.39 | **0.014** |
| Root Fe (mg g^-1^) | 1.51 | 0.220 | 0.02 | 0.977 | 0.07 | 0.935 |

**Table S3 Comparison of the alpha-diversity of rhizobia in root nodule pools of rooibos plants with high, medium and low root system biomass.** DNA sequence data for the *gyrB* gene was used to determine operational taxonomic units at 99% sequence identity, whose relative abundances were used to calculate the Simpson’s diversity index. The means with associated standard deviations are shown together with results of Tukey’s HSD mean comparison tests at p < 0.05.

| *Root biomass group* | *Richness* | | *Simpson Diversity* | |
| --- | --- | --- | --- | --- |
|  | *Mean (± SD)* | *Significance (Tukey’s HSD)* | *Mean (± SD)* | *Significance (Tukey’s HSD)* |
| High | 52.13 **±** 25.10 | a | 3.80 **±** 2.01 | a |
| Medium | 41.27 **±** 18.05 | b | 2.78 **±** 1.89 | a |
| Low | 30.79 **±** 14.92 | c | 2.49 **±** 1.53 | b |

| *ZOTU* | *Genus* | *Relative abundance (%)* |
| --- | --- | --- |
| ZOTU67 | *Mesorhizobium* | 17.56 |
| ZOTU131 | *Rhizobium* | 5.37 |
| ZOTU429 | *Brevundimonas* | 3.29 |
| ZOTU439 | *Bradyrhizobium* | 2.52 |
| ZOTU344 | *Bradyrhizobium* | 2.13 |
| ZOTU367 | *Herbaspirillum* | 1.54 |
| ZOTU221 | *Rhizobium* | 1.22 |
| ZOTU539 | *Bradyrhizobium* | 1.11 |
| ZOTU442 | *Bradyrhizobium* | 1.00 |
| ZOTU563 | *Rhizobium* | 0.53 |

**Table S4 Ten operational taxonomic units (OTU) unique to eight-month-old rooibos plants that were fertilized with sheep manure.** ZOTUs and their genus-affiliations are based on a gyrB DNA sequence data and relative abundances of each are provided.

**Table S5 Correlation between selected parameters of eight-month-old rooibos plants and the community composition and structure of the rhizobial root nodule symbionts as determined by a *gyrB* and *nodA* DNA sequencing data.** The parameters were chosen according to their expected relevance to describe rooibos above and belowground resource allocation. Mantel tests were run on the Euclidean distances of the plant parameters and the Bray-Curtis distances of the rhizobial root nodule community (n=258).

| *Plant parameter* | *gyrB* | | *nodA* | |
| --- | --- | --- | --- | --- |
|  | *Mantel r* | *P-value* | *Mantel r* | *P-value* |
| Plant dry biomass | 0.041 | 0.083 | -0.017 | 0.844 |
| Above:belowground dry biomass | 0.056 | 0.077 | 0.047 | **0.018** |
| # Nodules per plant | 0.052 | 0.066 | 0.002 | 0.455 |
| Leaf ^5^N | 0.010 | 0.341 | 0.031 | **0.049** |
| Leaf P conc. | 0.036 | 0.107 | 0.008 | 0.315 |
| Leaf N conc. | 0.029 | 0.159 | 0.004 | 0.407 |
| Leaf Mn conc. | -0.042 | 0.895 | 0.034 | **0.036** |
| Leaf Cu conc. | -0.044 | 0.922 | 0.036 | **0.031** |
| Root P conc. | 0.009 | 0.383 | 0.044 | **0.018** |
| Root Ca conc. | 0.051 | **0.036** | 0.041 | **0.008** |
| Root K conc. | -0.013 | 0.638 | 0.034 | **0.024** |
| Root Mg conc. | 0.041 | 0.083 | -0.017 | 0.844 |
